# Bruno 1 regulates cytoskeleton dynamics and a temporal splicing transition to promote myofibril assembly, growth and maturation in *Drosophila* flight muscle

**DOI:** 10.1101/2023.06.24.546382

**Authors:** Elena Nikonova, Marc Canela Grimau, Christiane Barz, Alexandra Esser, Jessica Bouterwek, Akanksha Roy, Heidemarie Gensler, Martin Heß, Tobias Straub, Ignasi Forne, Maria L. Spletter

## Abstract

Muscles undergo developmental transitions in gene expression and alternative splicing that are necessary to refine sarcomere structure and contractility. CUG-BP and ETR-3-like (CELF) family RNA binding proteins are important regulators of RNA processing during myogenesis that are misregulated in diseases such as myotonic dystrophy (DM1). Here we report a conserved function for Bruno 1 (Bru1, Arrest), a CELF1/2 family homolog in *Drosophila*, during early muscle myogenesis. Loss of Bru1 in flight muscles results in disorganization of the actin cytoskeleton leading to aberrant myofiber compaction and defects in pre-myofibril formation. Temporally-restricted rescue and RNAi knockdown demonstrate that early cytoskeletal defects interfere with subsequent steps in sarcomere growth and maturation. Early defects are distinct from a later requirement for *bru1* to regulate sarcomere assembly dynamics during myofiber maturation. We identify an imbalance in growth in sarcomere length and width during later stages of development as the mechanism driving abnormal radial growth, myofibril fusion and the formation of hollow myofibrils in *bru1* mutant muscle. Molecularly, we characterize a genome-wide transition from immature to mature sarcomere gene isoform expression in flight muscle development that is blocked in *bru1* mutants. We further demonstrate that temporally restricted Bru1 rescue can partially alleviate hypercontraction in late pupal and adult stages, but it cannot restore myofiber function or correct structural deficits. Our results reveal the conserved nature of CELF function in regulating cytoskeletal dynamics in muscle development, and demonstrate that defective RNA processing due to misexpression of CELF proteins causes wide-reaching structural defects and progressive malfunction of affected muscles that cannot be rescued by late-stage gene replacement.

## Introduction

Alternative splicing plays a key role in shaping the diverse contractile and morphological characteristics of different striated muscle fiber types (1,2). For example, the heart expresses short splice isoforms of Titin which contribute to the high passive resting stiffness of cardiomyocytes (3,4), while skeletal muscles with a lower passive resting stiffness express longer and more flexible Titin isoforms (5,6). Fast and slow muscle fibers express different isoforms of Troponin I (TnI) and Troponin T (TnT), resulting in differences in Ca^2+^ sensitivity and contractile dynamics (7). The fiber-type specific expression patterns of hundreds of exons are established during development, with transitions to mature isoforms promoting acquisition of fiber-type characteristic contractile properties (8–10). Although the functional differences between most splice isoforms are still unknown, misregulation of alternative splicing and isoform expression in muscle diseases such as dilated cardiomyopathies and myotonic dystrophies contributes to contractile dysfunction (11–13), highlighting the importance of RNA regulation to normal muscle function. Even different muscle fiber types in model organisms such as *Drosophila melanogaster* have distinct alternative splicing profiles (14,15), indicating that the regulation of alternative splicing and structural isoform expression plays a conserved role in fine-tuning muscle structure and contractile properties.

CUG-BP- and ETR-3-like factor (CELF) family RNA binding proteins (also known as Bruno-like proteins) are important regulators of RNA processing. CELF proteins contain three highly conserved RNA recognition motif domains (RRMs) that are jointly involved in binding to GU-rich recognition elements in RNA (16,17). They regulate diverse steps in RNA processing, from alternative splicing to mRNA trafficking, stability, decay and translation (18–20). In striated muscles, CELF proteins are involved in regulating developmental transitions in alternative splicing. CELF1 and CELF2 promote embryonic splicing patterns in vertebrate heart and skeletal muscle (21,22), for example promoting inclusion of cardiac troponin T (cTNT) exon 5 in embryonic heart affecting calcium sensitivity and contractility in mouse and chicken (23). CELF1/2 levels are downregulated 10-fold as heart and skeletal muscle mature (21,22), and overexpression of CELF1 during mouse heart development affects nearly 30% of developmental-associated splicing changes, largely promoting reversion to the embryonic splicing pattern (24). While CELF1/2 are downregulated in muscle development, Muscleblind-like family proteins MBNL1 and MBNL2 are in contrast upregulated and promote mature splicing and polyadenylation patterns (21,25,26). CELF1/2 and MBNL1/2 antagonistically co-regulate the alternative splicing of hundreds of exons in developing muscle (24,27). The physiological relevance of this regulatory interaction is illustrated by the severity of muscle phenotypes in myotonic dystrophy (DM) patients, where sequestration of MBNL1 through binding to a repeat expansion in the *DMPK* gene results in PKC-mediated stabilization and increased expression of the CELF1 protein (28,29), and a corresponding reversion from mature to embryonic isoform expression patterns (21,25,28,30). Thus, CELF proteins are a key component of the RNA regulatory network that defines muscle structure and contractile ability during myogenesis.

The conservation of CELF protein function in myogenesis provides an opportunity to explore foundational mechanisms of RNA regulation in muscle in greater detail. In zebrafish, CELF proteins are expressed in the developing mesoderm and Celf1 regulates somite development, binds to URE elements and can mediate splicing of a rat α-actinin mini-gene (31–33). In *C. elegans*, ETR-1, a CELF1 homolog, promotes muscle development through the regulation of alternative splicing and alternative 3’ exons (34,35). We and others have previously shown that Bruno 1 (Bru1, Arrest), a CELF1/2 family homolog in *Drosophila*, acts as a splicing factor during maturation of the indirect flight muscles (IFMs) to regulate growth in sarcomere length and myosin contractility (14,36). Bru1 expression is activated by the master regulator of the fibrillar muscle fate Spalt major (Salm), and hundreds of IFM-specific splice events in structural genes are lost after *bru1* RNAi knock-down (14,36). While the direct targets of Bru1 and detailed molecular mechanisms that contribute to the Bru1 phenotype are not known, loss of a Bru1-regulated, IFM-specific isoform of Stretchin-Myosin light chain kinase (Strn-Mlck) is sufficient to induce hypercontraction, short sarcomeres and loss of myofibers (14,37). Bru1 genetically interacts with RNA-binding protein Rbfox1 in IFM, resulting in complete loss of sarcomeric structure when both proteins are knocked-down and mirroring a regulatory interaction observed in mammals (27,38). Although Bru1 levels peak early in IFM development and Bru1 is downregulated in adult flies (38), all reported phenotypes for Bru1 affect later steps in sarcomere maturation after 48 h APF (14,36,38,39). The question therefore arises whether Bru1 has a function during early stages of IFM formation, congruent with the role of CELF1/2 in fetal muscle in vertebrates, or if CELF function in *Drosophila* muscle is mechanistically distinct.

The *Drosophila* IFMs are an established and disease-relevant model for exploring basic mechanisms of muscle development and sarcomere assembly. Sarcomere structure is conserved, and in both insects and vertebrates, sarcomeres are built of actin thin filaments anchored at the Z-disc, myosin thick filaments anchored at the M-line, and Titin connecting filaments that span the thin and thick filaments (40–42). Myosin binding to actin and filament sliding provides contractile force, while Titin influences muscle stiffness, force generation and sarcomere length (43–46). Analysis of *Drosophila* models of human disease, for example myotonic dystrophy, X-linked centronuclear myopathy, nemaline myopathy and Duchenne muscular dystrophy, have proven informative and offer relevant insight into disease pathology (13,47,48). Work in *Drosophila* also provides insight into conserved developmental mechanisms of myogenesis, including myoblast fusion, tendon attachment, sarcomerogenesis, growth and myofibril maturation (49–51). The *Drosophila* IFMs consist of six dorsal-longitudinal (DLM) and seven dorsal-ventral myofibers (DVM) in each hemisphere (52,53). IFM myoblasts proliferate on the wing-disc hinge, and then migrate and fuse to form IFM myotubes (54,55). IFM myotubes establish tendon connections around 16-20 hours after puparium formation (h APF) (56), and then compact and undergo myofibrillogenesis around 32 h APF (42,57,58). Sarcomeres are added to myofibrils as myofibers grow dramatically in length to span the entire thorax by 48 h APF, and from 60 h to 90 h APF sarcomeres grow to their mature size of 3.2 μm in length and 1.2 μm in width (42,59,60). After 48 h APF, myofibrils undergo a maturation process where a switch in gene expression facilitates establishment of asynchronous and stretch-activation properties of fibrillar IFM (37,61). This detailed understanding of myogenesis in a conserved genetic model system is a powerful tool that can be applied to understand how RNA regulation impacts sarcomere assembly and maturation.

Here we report that the early requirement for CELF protein function in myogenesis is conserved in *Drosophila* Bruno 1 (Bru1). We generated a novel CRISPR-mediated mutant in *bru1* that revealed early phenotypes in cytoskeletal organization. Temporally-restricted rescue and *bru1* RNAi knockdown demonstrated how initial cytoskeletal defects are propagated and disrupt later steps in sarcomere growth and maturation. We further define a later requirement during myofiber maturation for *bru1* to regulate sarcomere assembly dynamics, where abnormal radial growth in *bru1* mutant muscle promotes myofibril fusion and the formation of hollow myofibrils. Our data moreover identify a previously uncharacterized genome-wide transition from immature to mature sarcomere gene isoform expression in IFMs that is blocked in *bru1* mutants. Consistent with the pleiotropic nature of the CELF misregulation phenotype, temporally restricted expression of Bru1 cannot restore myofiber function or correct structural deficits, but does partially alleviate adult-stage hypercontraction. Our results reveal a conserved role for CELF family proteins to fine-tune sarcomere structure and function, and identify multiple distinct developmental mechanisms that contribute to the *bru1* mutant phenotype in IFM.

## Results

To investigate the function of Bru1 in *Drosophila* IFM development, we generated a new CRISPR allele that we refer to as *bru1^M3^*. *bru1^M3^* is a truncation allele resulting from the integration of a splice-trap cassette upstream of *bru1* exon 18 (Fig S1 A, B) that results in a near complete loss of detectable *bru1* mRNA and protein expression (Fig S7 A, C, C’). The splicing of remaining *bru1* transcripts is redirected into the splice acceptor of the cassette instead of into exon 18, generating an early termination that effectively deletes the most C-terminal 88 amino acids in RRM3 of all *bru1* isoforms (Fig S1 C, D). Based on phenotypes reported for the *aret^QB72^* allele (EMS-induced stop at position 404) (14,62,63), as well as point mutations in RRM3 at positions 521 and 523 (64), *bru1^M3^* is predicted to be a phenotypic null allele. Like other *bru1* alleles (62,63), *bru1^M3^* is male and female sterile. Consistent with the reported IFM phenotype of *bru1* RNAi knockdown (*bru1-IR*) in adult flies (14,36), we found that *bru1^M3^* mutants are flightless and display a loss of myofibril and sarcomere architecture (Fig 1 A, B). We further confirmed the specificity of this phenotype over deficiency Df(2L)BSC407, which covers the *bru1* locus (Fig 1 B, Fig S1 E, E’, F, F’, G). Together, these findings validate the nature and specificity of the *bru1^M3^* allele, and provide independent confirmation of a function for Bru1 in IFM development.

**Fig 1.**
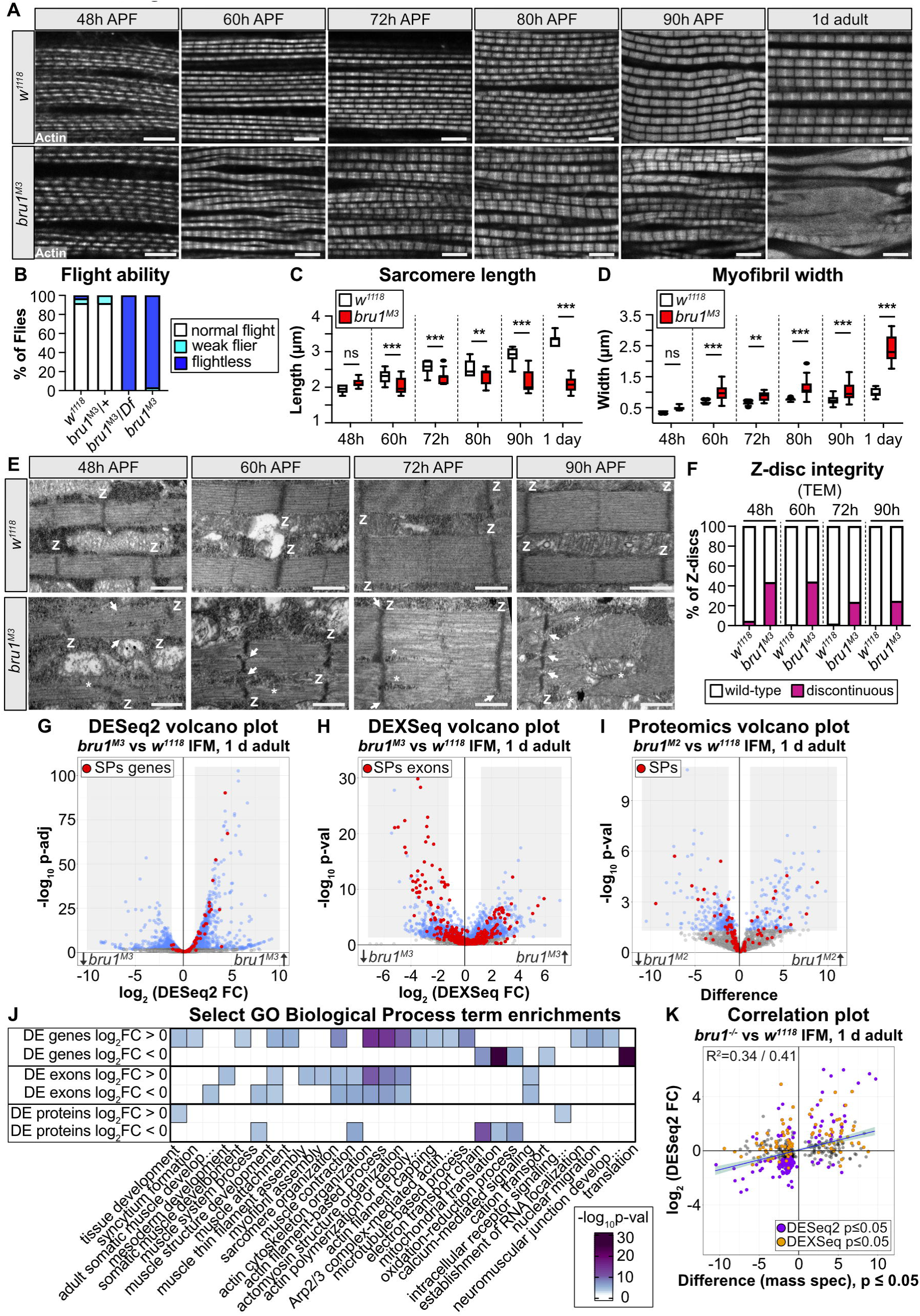
*bru1* mutant flight muscle displays misregulated sarcomere protein expression and progressively severe phenotypes during myofibril maturation. **(A)** Single-plane confocal images from thorax hemi-sections of *w^1118^* and *bru1^M3^* at 48 hour (h), 60 h, 72 h, 80 h, 90 h after puparium formation (APF) and 1 d adult flies. Phalloidin stained actin, grey; scale bar = 5 μm. **(B)** Quantification of flight ability. N > 30 flies for each genotype. **(C-D)** Quantification of the sarcomere length (C) and myofibril width (D) from (A). Boxplots are shown with Tukey whiskers, outlier data points marked as black dots. Significance determined by ANOVA and post hoc Tukey (ns, not significant; **, p < 0.01; ***, p < 0.001). **(E)** Transmission electron microscopy (TEM) images of *w^1118^* and *bru1^M3^* sarcomere ultrastructure at 48 h, 60 h, 72 h and 90 h APF. Defects in *bru1^M3^* are already apparent at 48h APF. Z-discs, “Z”; myofibril splitting and discontinuous Z-discs, white arrows; cytoplasm or mitochondrial inclusions, white asterisks; scale bar = 1 μm. **(F)** Quantification of Z-disc integrity in (E). N > 20 single planes for each individual genotype and time point. **(G-H)** mRNA-Seq volcano plots of DESeq2 gene expression (G) and DEXSeq exon use (H) changes in 1 d adult *bru1^M3-/-^*versus *w^1118^* IFM. Sarcomere proteins (SPs) are notably affected (red dots). Grey boxes denote a threshold of abs(log_2_ fold-change) ≥ 1and p ≤ 0.05, with significant events coloured blue. **(I)** Volcano plot of peptide group expression (I) changes in *bru1^-/-^* IFM from 1 d adults. Grey boxes denote a threshold of abs(Difference) ≥ 1 and p ≤ 0.05, significant peptides are colored blue. **(J)** Heatmap of select significantly enriched biological process GO terms in the differentially expressed (DE) genes, exons and proteins. **(K)** Dot plot of the correlation between significantly DE peptide groups and their corresponding mRNA expression level in *bru1^-/-^* versus w^1118^ IFM. Proteins with a significantly DE exon (DEXSeq) are colored orange, and those significantly DE at the gene level (DESeq2) are colored purple. The Pearson’s / Spearman’s correlation coefficients (top left corner) and regression line (blue) indicate a weak but positive correlation.

### Growth in sarcomere width and length is imbalanced in *bru1^M3^* mutant IFM

We used the *bru1^M3^* allele to perform a detailed analysis of IFM development from 48 h after puparium formation (APF) through 1 d adult in thorax hemi-section preparations. We reasoned that a phenotypic null mutant might produce a stronger phenotype than the *bru1-IR* knockdown used in previous studies (14,36). *bru1^M3^* mutant myofibrils are severely degraded in 1 d adult, but are structurally intact until 90 h APF. However, already at 60 h APF mutant myofibrils are split and irregular, and neighboring myofibrils appear partially aligned (Fig 1 A). From 48 to 90 h APF, sarcomeres in control *w^1118^* flies grow significantly in length from 2.0 ± 0.1 to 2.9 ± 0.2 μm (ANOVA, p < 0.001), while sarcomeres in *bru1^M3^* show an insignificant change from 2.1 ± 0.1 to 2.2 ± 0.3 μm (ANOVA, p = 0.96) (Fig 1 A, C). This indicates that growth in sarcomere length in *bru1^M3^* IFM is arrested after 48 h APF. By contrast, already at 60 h APF in *bru1^M3^*, myofibrils are significantly wider than in wildtype (0.99 ± 0.2 versus 0.66 ± 0.04 μm, ANOVA, p < 0.001), and this difference is maintained throughout late pupal stages (Fig 1 D). Myofibrils in *bru1^M3^*IFM significantly increase in width from 0.48 ± 0.06 to 1.03 ± 0.26 μm (ANOVA, p < 0.001) from 48 h to 90 h APF (Fig 1 D). This suggests that although *bru1^M3^* sarcomeres do not grow in length after 48 h APF, they do grow in width.

To better understand the myofibril growth defects in *bru1^M3^* mutant IFMs between 48 h to 90 h APF, we performed an ultrastructural time course analysis using transmission electron microscopy (TEM). Sarcomeres in *bru1^M3^* IFM were significantly shorter and thicker than *w^1118^* sarcomeres at 60 h, 72 h and 90 h APF (Fig 1 E, Fig S1 H, I), confirming our observations from thorax hemi-sections (Fig 1 A, C, D). Notably, already at 48 h APF, split myofibrils as well as discontinuous and misaligned z-discs were evident in *bru1^M3^* IFMs (Fig 1 E, F, Fig S1 J, K), indicating that myofibril defects in *bru1* mutants arise earlier than previously reported (14,36). Myofibril splitting and z-disc defects were also detected in *bru1^M3^* at the 60 h, 72 h and 90 h timepoints (Fig 1 E, F, Fig S1 J, K). Peculiarly, starting from 60 h APF, we could identify cytoplasmic components and even mitochondria trapped in the middle of otherwise continuous myofibrils (Fig 1 E), suggestive of an underlying defect in radial growth or myofibril fusion. This data thus reveals two novel aspects of the *bru1^M3^* phenotype in IFM that we explore in more detail below: 1) during myofibril maturation there is an imbalance between growth in sarcomere length and width, and 2) defects in *bru1^M3^*mutant myofibril structure are already evident at 48 h APF.

### Misregulation of gene expression and splicing lead to protein expression defects in *bru1^M3^* muscle

We next investigated the molecular phenotype underlying the myofibril defects observed in *bru1^M3^* IFM. We performed mRNA-Seq and whole proteome mass spectrometry on IFM dissected from 1 d adult wildtype *w^1118^* or *bru1* mutant flies, to evaluate changes on both the RNA and protein levels (Fig 1 G, H, I, Table S1). A differential expression analysis with DESeq2 revealed hundreds of significant changes in gene expression in the mRNA-Seq data (Fig 1 G). Upregulated genes were enriched for biological process gene ontology (GO) terms such as “muscle attachment,” “sarcomere organization,” “actin cytoskeletal organization,” “actin filament capping,” and “establishment of RNA localization” (Fig 1 J). Downregulated genes were in contrast enriched for terms such as “translation,” “cation transport,” and “oxidation-reduction process.” Using DEXSeq, we further detected hundreds of significant changes in exon use in *bru1^M3^* versus *w^1118^* IFM (Fig 1 H), reflecting changes in alternative splicing as well as alternative promoter use. Notably, both upregulated and downregulated exons were enriched for GO terms such as “sarcomere organization,” “actin cytoskeleton organization,” “muscle contraction,” and “calcium-mediated signaling” (Fig 1 J). This likely reflects isoform switches in structural genes, as for example sarcomere proteins (SPs) display both up- and downregulated exons (Fig 1 H), which we investigate in more detail below. Interestingly, on the gene level, SPs are mostly upregulated in *bru1^M3^*versus *w^1118^* IFM, potentially reflecting transcriptional compensation in response to changes in isoform use (Fig 1 G). We conclude that loss of Bru1 function leads to changes in both gene expression and alternative splicing.

We complimented our transcriptomic data with a proteome analysis from 1 d adult IFM, to evaluate if mRNA-level changes translate to altered protein expression. We grouped detected peptides into protein groups, such that peptides from the same gene that are unique to different protein isoforms form distinct protein groups. Analysis of the proteomics data revealed significant changes in the expression of hundreds of protein groups, with a bias towards downregulation (Fig 1 I). Downregulated proteins were enriched in GO terms such as “muscle system process,” “muscle contraction,” “electron transport chain,” and “oxidation-reduction process,” while upregulated proteins were enriched in “tissue development” and “intracellular receptor signaling process” (Fig 1 J). As in the exon use analysis, we see up- and downregulation of different sets of protein groups from sarcomere proteins (Fig 1 I). We then tested if there is a relationship between the changes in gene expression at the mRNA and protein level, and observed a weak but positive correlation for all significantly changed protein groups (Pearson’s R^2^ = 0.34, Spearman’s R^2^ = 0.41) (Fig 1 K). Interestingly, we saw that both changes in gene expression as well as changes in exon use correlated with differential expression on the protein level (Fig 1 K, Fig S1 L). We also noted that different categories of genes show distinct patterns of regulation on the RNA and protein level. Cytoskeletal genes such as SPs, genes involved in actin cytoskeleton organization, and microtubule-associated genes tend to be upregulated at the gene level, but show both up- and downregulation in the DEXSeq and proteomics data (Fig S2 A). Mitochondrial and fibrillar core genes (14), by contrast, have up- and downregulated exons, but are downregulated on both the gene and protein level (Fig S2 A). We conclude that Bru1-mediated changes to the IFM transcriptome are complex and indeed alter protein and protein isoform expression.

### Protein-level misexpression reflects alternative splicing changes in *bru1^M3^* IFM

Reasoning that the strong changes in structural gene expression at both the mRNA and protein level might contribute to the myofibril defects in *bru1^M3^* IFM, we analyzed sarcomere protein expression in greater detail. We noted that while 39% (9 of 23) of significantly differentially expressed sarcomere proteins (SPs) were differentially regulated on the mRNA level (compared to 36% (184 of 514) of all significantly differentially expressed proteins), 74% (17 of 23) displayed significantly differential exon use (compared to 84 of 514 or 16% of all proteins) (Fig 2 A, Fig S2 A). We therefore compared changes in gene expression, exon use and protein isoform expression in SPs (Fig 2 B, Table S2). We found a clear correspondence between changes in mRNA expression or splicing and altered protein expression level. For some SPs, for example Fhos, Ilk or Mlp84B, gene expression changes match observed protein level changes (Fig 2 B). For other SPs, for example Mlp60A, wupA, Mhc, bt and Tm1, we see a clear switch in isoform expression, where changes in exon use match observed protein isoform changes (Fig 2 B). This data also illustrates the breadth of sarcomere assembly processes impacted in *bru1* mutant muscle, as misexpressed SPs include previously identified regulators of thin and thick filament growth, structural components of the z-disc and M-line, components of the integrin and cell adhesion machinery, and regulators of actomyosin interactions (65).

**Fig 2.**
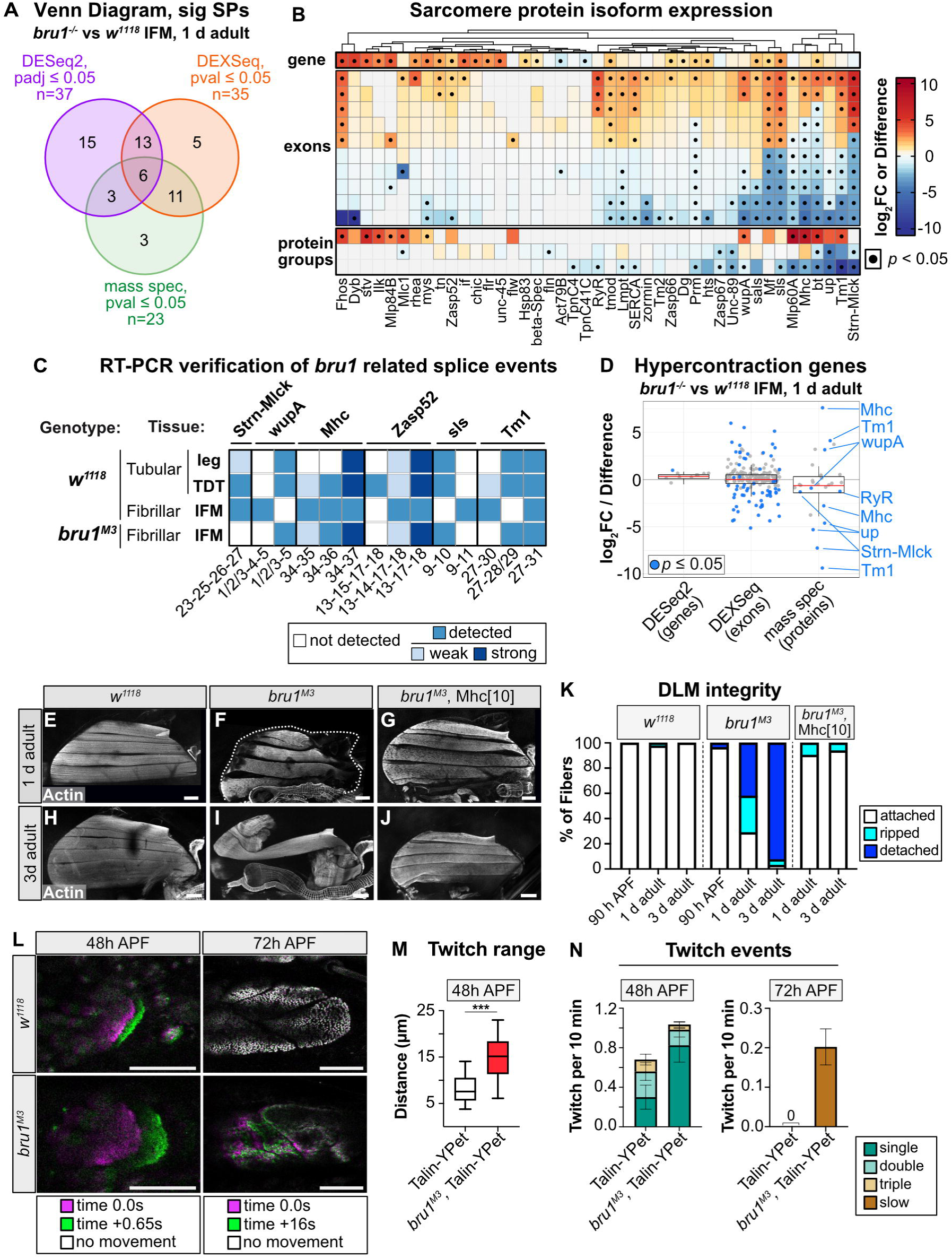
Misexpression of muscle-type specific protein isoforms results in hypercontraction and abnormal contractile dynamics in *bru1^-/-^* IFM. **(A)** Venn diagram of the overlap between SPs differentially expressed at the gene level (purple), the protein level (green) or that have differentially used exons (orange). **(B)** Heirarchical clustering and heat map of the coordinated changes in SP gene expression (top, DESeq2 log_2_FC), exon use (middle, top 10 DEXSeq log_2_FC values) and protein expression (bottom, Perseus Difference of the top 3 protein groups). Black dot denotes p ≤ 0.05. **(C)** Summary heatmap of RT-PCR results (Fig S2) verifying muscle-type specificity and loss or gain of specific SP alternative splice events in *bru1^-/-^* IFM in *Strn-Mlck*, *wupA*, *Mhc*, *Zasp52*, *sls* and *Tm1* (not detected by PCR, white; detected by PCR, blue; weak band, light blue; strong band, dark blue). **(D)** Boxplot of gene, exon and protein level expression changes in genes with associated hypercontraction phenotypes (Table S1, Table S2). Significantly DE proteins in *bru1^M3^* IFM are labelled; blue dot denotes p ≤ 0.05. **(E-J)** Confocal Z-stack images of DLM myofiber in w*^1118^*, *bru1^M3^* and *bru1^M3^*, Mhc[10] from 1 d and 3 d adults. Myofiber loss and hypercontraction in *bru1^-/-^*IFM is alleviated in the Mhc[10] background (G, J). Thorax boundaries in (F), dashed line; phalloidin stained actin, grey; Scale bar = 100 μm. **(K)** Quantification of myofiber tearing and detachment phenotypes from (E-J) at 90 h APF, 1 d and 3 d adults. N > 40 myofibers from at least 10 flies for each individual genotype and time point. **(L)** Snapshots from live movies of talin-YPet labeled DLMs at 48 h and 72 h APF from *w^1118^*and *bru1^M3^* animals. Time 0.0 (magenta) is overlaid with time +0.65 s (green; at 48 h APF) or +16 s (green; at 72 h APF). A complete overlap (white) depicts no movement. Scale bar = 50 μm. **(M)** Quantification of distance of maximum myofiber extension at 48 hr APF. Boxplots are shown with Tukey whiskers, significance by unpaired t-test (***, p < 0.001). **(N)** Quantification of spontaneous contraction events per fiber per 10 min in control and *bru1^M3^*. At 72 h APF, *bru1^M3^* DLM fibers continue to undergo slow, unidirectional extension. N > 50 fibers/10 animals for each genotype and time point. Error bars = SEM.

To independently confirm changes in alternative splicing and protein isoform expression in *bru1^M3^* IFM, we used GFP-tagged reporters and RT-PCR to test a panel of events in *Strn-Mlck*, *wupA*, *Mhc*, *Zasp52*, *sls* and *Tm1* (Fig 2 C). We confirmed the loss of expression of *Strn-Mlck* isoform R (exons 23 and 25) at both the RT-PCR and protein level in *bru1^M3^* IFM (Fig 2 C, Fig S2 C, F, G, G’, H, H’), which was previously shown to result in IFM hypercontraction (14). In *wupA*, which encodes Troponin I (66), we confirmed an isoform switch from the IFM- to the tubular-specific termination (exon 4) at both the mRNA and protein levels (Fig 2 C, Fig S2 D, I, J, J’, K, K’). We found that an alternative termination of *Mhc* that is used at early stages of IFM development and in tubular muscle (exon 37) (39,61) is expressed at increased levels in 1 d adult *bru1^M3^* IFM at both mRNA and protein levels (Fig 2 C, Fig S2 E, L, M, M’, N, N’). Interestingly, this Mhc isoform is incorporated uniformly across the width of the myofibril in *bru1^M3^*, instead of restricted to the center of the myofibrils as observed in control IFM (Fig S2 M, M’, N, N’). We further could confirm a switch in exon use in *Zasp52* exon 14/15, *sls* exon 10 and *Tm1* exon 28/29 and 30 (Fig 2 C, Fig S2 O, P, Q). This data validates our transcriptome and proteome analyses, and confirms that Bru1 regulates alternative splicing and protein isoform expression of sarcomere proteins in IFM.

### *bru1^M3^* mutant myofibers experience abnormal contractility throughout development

Several of the proteins we verified to have altered isoform expression in *bru1^M3^* mutants share a function in regulating actomyosin interactions, suggesting that normal contractile dynamics may be altered after loss of Bru1. Specifically, when we curated a list of genes with reported hypercontraction phenotypes in fly muscle, we noticed significant changes in alternative splicing and protein isoform expression in Mhc, Tm1, wupA, RyR, Strn-Mlck and up (TnT) (Fig 2 D, Fig S2 G’, J’, M’). Misregulated actomyosin interactions can lead to hypercontraction, a condition characterized by aberrant contractility and short, thick sarcomeres (67–69). In our time course analysis, we noticed a dramatic increase in the severity of the *bru1^M3^* phenotype at late stages of pupal development (Fig 1 A, C, D). From 80 h APF to 1 d adult, sarcomeres in *bru1^M3^*shorten significantly from 2.3 ± 0.2 to 2.1 ± 0.2 μm (ANOVA, p = 0.009), while *w^1118^* sarcomeres grow from 2.5 ± 0.2 to 3.3 ± 0.1 μm, respectively (ANOVA, p < 0.001) (Fig 1 C). By contrast, myofibril width more than doubles in *bru1^M3^* mutants from 1.1 ± 0.3 μm at 80 h APF to 2.4 ± 0.4 μm in 1 d adult (ANOVA, p < 0.001), but only increases in wildtype from 0.76 ± 0.04 to 1.0 ± 0.1 μm (ANOVA, p = 0.002) (Fig 1 D). Additionally, while IFM myofibers are attached at 90 h APF in *bru1^M3^*mutants, at 1 day 71% and by 3 days 97% of myofibers are torn or completely detached (Fig 2 E, F, H, I, K). A hypercontraction phenotype can be rescued by minimizing actomyosin forces, and we confirmed that *bru1^M3^* myofibers remain attached in an *Mhc^10^* mutant background in both 1- and 3-day adults (Fig 2 G, J, K). This verifies that myofiber loss is indeed myosin-activity dependent, consistent with previous RNAi results (14,36). We conclude that *bru1^M3^*myofibrils experience hypercontraction from 80 h APF, characterized by progressive shortening and thickening of sarcomeres and eventual myofiber loss in adult flies.

Actomyosin-dependent tension also plays an important role during early stages of IFM development to organize and refine sarcomere structure (70,71). Spontaneous contractions, or twitches, are evident at 34 h APF shortly after sarcomere assembly, reach peak intensity around 48 h APF, and are suppressed by 72 h APF as IFM myofibrils develop stretch-activation properties (37). To determine if actomyosin contractility is disrupted in *bru1^M3^* mutants during IFM development, we performed live imaging of IFMs labeled with Talin-YPet. At 48 h APF, we could detect twitch events in both *bru1^M3^* and control *w^1118^* myofibers (Fig 2 L, Media File 1). Strikingly, contractions in *bru1^M3^*myofibers were stronger than in *w^1118^* myofibers, resulting in a greater displacement of the myofiber tip (Fig 2 M), and occurred more frequently (Fig 2 N). At 72 h APF, we were unable to detect any movement in *w^1118^* myofibers (Fig 2 L, N). Unexpectedly, in 17 of 89 *bru1^M3^*myofibers (19%), we observed a slow twitch movement (Fig 2 L, N, Media File 2). These fibers slowly contracted over a period of 16 seconds, as compared to less than one second at 48 h APF, and did not efficiently reextend. We conclude that actomyosin interactions are atypical throughout IFM development in *bru1* mutants, with abnormal contractility first contributing to sarcomere organization defects and then resulting in hypercontraction in late pupa and adult flies.

**Media File 1. Movie of spontaneous contractions at 48 h APF in wild-type and *bru1^M3^* DLM myofibers.**

Spontaneous contractions in control and mutant IFMs at 48 h APF. From left to right: wild-type DLM performing a single twitch; wild-type DLM performing a double twitch; wild-type DLM performing a triple twitch, *bru1^M3^* DLM performing a single twitch. Timer, bottom left; Scale bar = 50 μm.

**Media File 2. Movie of spontaneous contractions at 72 h APF in wild-type and *bru1^M3^* DLM myofibers.**

Wild-type control IFMs do not spontaneously contract at 72 h APF (left). A slow, one-directional contractile movement is observed in *bru1^M3^*myofibers at 72 hr APF (right). Timer, bottom left; Scale bar = 50 μm.

### Formation of hollow myofibrils is driven by misregulated radial growth and myofibril fusion

Another subset of sarcomere proteins misexpressed in *bru1^M3^*IFM, including *Fhos*, *sals*, *sls*, *Mlp84B*, *Unc-89*, *tmod* and *Zasp52*, are known regulators of sarcomere growth and integrity (65). These genes caught our attention, as we observed splitting and an abnormal balance between growth in length and width in *bru1^M3^* sarcomeres (Fig 1 A, E). We also noticed that not just SPs, but more broadly actin cytoskeleton and microtubule-associated genes, are misregulated in *bru1^M3^*(Fig 3 A, Fig S2 A). When we examined actin gene expression, we found altered expression ratios in our mRNA-Seq data that we could further confirm by RT-qPCR (Fig 3 B, C, Fig S3 A), including a dramatic upregulation of cardiac actin Act57B. Previous studies have shown that actin genes have differential abilities to integrate into the growing sarcomere, and misexpression of cardiac Act57B disrupts IFM function (72,73). We reasoned that the altered ratio in actin gene expression in *bru1^M3^* IFM, together with changes in key cytoskeletal regulators, might lead to aberrant sarcomere growth.

**Fig 3.**
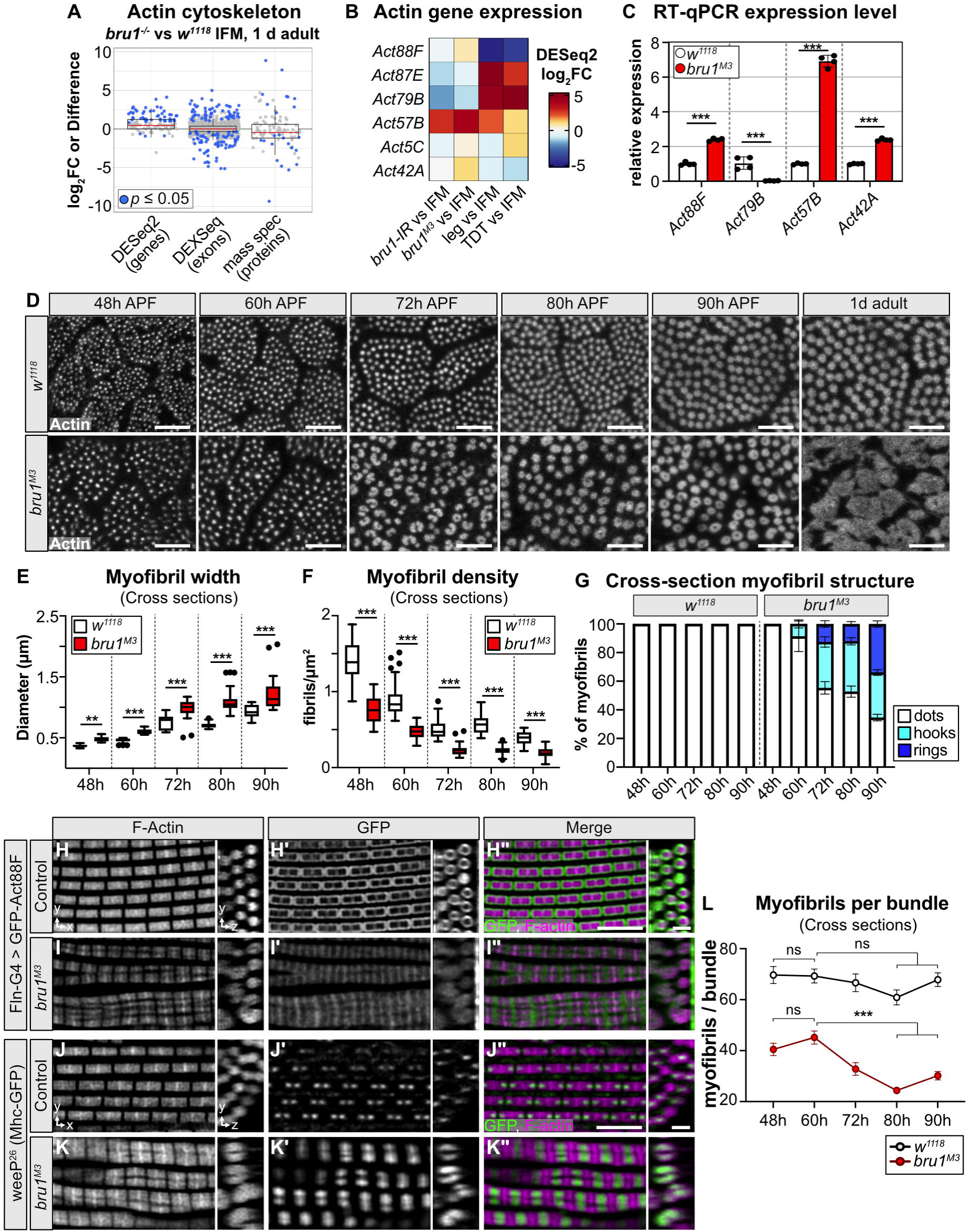
Hollow myofibril formation in *bru1* mutants is an active process resulting from defective expression and splicing of cytoskeletal genes and aberrant sarcomere growth. **(A)** Boxplot of gene, exon and protein level expression changes between 1 d adult *bru1^-/-^* and w^1118^ IFM in genes from GO term “actin cytoskeleton”. Blue dot denotes p ≤ 0.05. **(B)** Heatmap of *Actin* gene expression in *bru1-IR*, *bru1^M3^*, leg and TDT as compared to wildtype IFM. **(C)** RT-qPCR verification of *Actin* gene expression levels between *bru1^M3^* and wildtype *w^1118^* IFM. **(D)** Confocal micrographs of DLM myofibril cross-sections in *w^1118^* and *bru1^M3^*at 48 h, 60 h, 72 h, 80 h, 90 h APF and 1 d adult. Phalloidin stained actin, grey; Scale bar = 5 μm. **(E-F)** Quantification of myofibril width (E) and density (F) in (D). Boxplots are shown with Tukey whiskers, outlier data points marked as black dots. Significance determined by ANOVA with post-hoc Tukey (**, p < 0.01; ***, p < 0.001). **(G)** Quantification of myofibril structural morphology in (D). *bru1^M3^* myofibrils progressively form hook and ring structures starting from 60 h APF. N > 10 animals per genotype and time-point. Error bars = SEM. **(H-I”)** Deconvoluted confocal images at 90 h APF of *Fln*-Gal4 driven UASp-GFP-Actin88F incorporation into wild-type (H-H”) and *bru1^M3^* (I-I”) sarcomeres. Both xy- and zy-projections are shown. *Fln*-Gal4 expression from ∼56 h APF results in a box-like pattern of GFP-Act88F incorporation (H’-H”) into growing wildtype sarcomeres, which is abnormal in *bru1^M3^* (I’-I”). GFP, green; phalloidin stained actin, magenta; Scale bar = 5 μm. **(J-K”)** Deconvoluted confocal images at 90 hr APF of Mhc-weeP26-GFP incorporation into wild-type (J-J”) and *bru1^M3^* (K-K”) sarcomeres. Expression of GFP-labeled Mhc isoforms containing exon 37 is restricted to early developmental stages in IFM, resulting in a dot-like pattern flanking the M-line in wild-type sarcomeres (J’-J”) which is disrupted in *bru1^M3^*(K’-K”). weeP26-GFP, green; phalloidin stained actin, magenta; Scale bar = 5 μm. **(L)** Quantification of myofibril number per fiber bundle in *w^1118^*and *bru1^M3^* at 48 h, 60 h, 72 h, 80 h and 90 h APF. Plot represents the mean ± SEM. Significance determined by ANOVA and post-hoc Tukey (ns, not significant; **, p < 0.01). N > 8 animals per genotype and time-point.

To evaluate defects in lateral or radial sarcomere growth in greater detail, we performed a cross-section time course analysis of *bru1^M3^* IFM development. In wildtype *w^1118^* IFM, myofibrils are organized into distinct bundles and grow uniformly in the radial direction from 48 h to 1 d adult, maintaining a consistent, circular appearance in cross-section (Fig 3 D). Myofibrils in *bru1^M3^* flies are also organized into bundles, but already at 48 h APF, mutant myofibrils are significantly wider than in wildtype (0.48 ± 0.04 μm versus 0.37 ± 0.02 μm, respectively, ANOVA p = 0.001) (Fig 3 D, E), and there are significantly fewer myofibrils (0.76 ± 0.02 μm versus 1.4 ± 0.2 myofibrils per μm^2^, ANOVA p < 0.001) (Fig 3 F). Strikingly, although *bru1^M3^* myofibrils appear uniform and circular at 48 h APF, by 60 h APF there is variability among the sizes of individual myofibrils and some appear more oval or hook-like than circular (Fig 3 D, G). As the myofibrils develop from 60 h to 90 h APF, this hooked phenotype becomes more pronounced, affecting 8.7 ± 2.8% of myofibrils at 60 h APF and 31-35% of myofibrils at 72 h, 80h and 90h APF (Fig 3 G). Concurrently, we see the progressive development of rings, or hollow myofibrils, affecting 12.6 ± 2.2% of myofibrils at 72 h APF and 33.9 ± 2.2% of all myofibrils at 90 h APF (Fig 3 D, G). In 1 d adults, *bru1^M3^* myofibrils are highly irregular and display a strong atrophy phenotype. Taken together, this analysis reveals a distinct temporal progression in the appearance of myofibril defects, suggesting atypical and irregular radial growth during myofibril maturation. Moreover, such growth could provide a mechanism whereby cytoplasmic components or even organelles become trapped in the middle of myofibrils, as we observed in our TEM data (Fig 1 E).

To test if sarcomere growth is indeed aberrant in *bru1^M3^*IFM, we adapted an approach to visualize thin filament growth dynamics using temporally-restricted expression of a GFP-tagged actin (60). We expressed UAS-GFP-Act88F starting around 56 h APF using Flightin-Gal4 (Fln-G4), which allowed us to monitor actin incorporation over the critical time period for hollow myofibril formation. In control sarcomeres from 90 h APF flies, we observed a box-like labeling pattern, reflecting GFP-Act88F integration both laterally as well as at the plus and minus ends of the thin-filament (Fig 3 H, H’, H’’, Fig S3 B-B’’’, Media File 3). In *bru1^M3^* mutants, this pattern is significantly altered. GFP-Act88F is incorporated weakly at the z-disc and strongly and uniformly across the M-line (Fig 3 I, I’, I’’, Fig S3 C-C’’’, Media File 4). This is striking, as sarcomeres do not grow appreciably in length in *bru1^M3^* mutants, thus indicating a defect either in thin-filament stability or capping protein function. Moreover, the strong lateral incorporation of GFP-Act88F coupled with exclusion from the central core of the sarcomere observed in control flies is not evident in *bru1^M3^*(Fig 3 H’’, I’’), even though mutant sarcomeres exhibit excessive radial growth from 60 h to 90 h APF (Fig 3 D, E). We interpret this to reflect loose packing of the thin-filament lattice allowing actin integration across the sarcomere, as well as radial integration of other unlabeled actin isoforms. These data reveal a clear defect in thin filament growth dynamics, and identify separable phenotypes in processes controlling growth in sarcomere length and width.

**Media File 3. Movie of a 3D reconstruction of GFP-Act88F integration into wild-type sarcomeres.**

Movie of an x,y,z-axis reconstruction from a confocal image of Fln-Gal4 driving UASp-GFP-Act88F in a control *w^1118^* background. GFP labelling is restricted to a box-like pattern marking radial thin-filament addition and actin integration into thin filaments after 56 h APF. GFP, green; phalloidin stained actin, red.

**Media File 4. Movie of a 3D reconstruction of GFP-Act88F integration into *bru1^M3^* mutant sarcomeres.**

Movie of an x,y,z-axis reconstruction from a confocal image of Fln-Gal4 driving UASp-GFP-Act88F in a *bru1^M3^* background. GFP labelling is altered in comparison to wild-type, revealing a lack of GFP-Act88F integration into thin-filaments added radially after 56 h APF and abnormal actin integration across the entire thin filament with enrichment at the z-disc and M-line. GFP, green; phalloidin stained actin, red.

We next evaluated the localization of Mhc^Wee-P26^-GFP, to test if thick filament growth is also disrupted in *bru1^M3^* myofibrils. In control sarcomeres, Mhc^Wee-P26^-GFP is localized in two central dots flanking the M-line (Fig 3 J, J’, J’’, Fig S3 D, E-E’’’’, Media File 5), because a developmental switch in Mhc isoform expression results in incorporation of an unlabeled Mhc isoform after 48 h APF (61). This isoform switch is partially impaired in *bru1^M3^* IFM, so Mhc^Wee-P26^-GFP is continuously expressed (39), resulting in GFP incorporation across the thick filament (Fig 3 K, K’, K’’, Fig S3 F). Interestingly, Mhc^Wee-P26^-GFP can be seen to form an irregular and asymmetric hook-like pattern at the M-line in *bru1^M3^* sarcomeres (Fig 3 K#x2019;’, Fig S3 F, G-G’’’’, Media File 6), indicative of M-line misalignment or instability. We also observed myofibrils with two or more seemingly distinct Mhc^Wee-P26^-GFP dots (Fig 3 K’’, Media File 6), suggesting that large, irregular myofibrils may result from myofibril fusion. Logically, neighboring myofibrils undergoing abnormal radial growth might grow into one another and fuse, contributing to the formation of hollow myofibril structures. To test this, we calculated the number of myofibrils per bundle in our cross-section data (Fig 3 D), and found a significant reduction from 60 h to 80 h APF in *bru1^M3^* (45 ± 9 to 24 ± 5 myofibrils per bundle, ANOVA, p < 0.001) but not in *w^1118^* IFM (69 ± 15 to 68 ± 14 myofibrils per bundle, ANOVA, p = 0.30) (Fig 3 L). Interestingly, this partial fusion of myofibrils does not reflect a switch in the fiber-type fate of the IFMs, as the majority of tubular genes are not misexpressed in *bru1^M3^* IFM (Fig S3 H). Similarly, while there is misuse of tubular exons in *bru1^M3^* IFM, many tubular exons are not strongly affected (Fig S3 H), indicating that loss of Bru1 does not result in a complete switch in fiber-type specific splicing, but rather loss of IFM-specific splice events. Taken together, our data support a mechanism whereby altered expression ratios of actins and cytoskeletal regulators impairs sarcomere growth in length, but concurrently promotes aberrant radial growth leading to myofibril fusion and the formation of hollow myofibrils during myofibril maturation in *bru1^M3^* IFMs.

**Media File 5. Movie of a 3D reconstruction of Mhc^Wee-P26^-GFP integration into wild-type sarcomeres.**

Movie of an x,y,z-axis reconstruction from a confocal image of Mhc-weeP26-GFP. GFP signal is restricted to two dots flanking the M-line, reflecting the first bipolar myosin filaments assembled into pre-myofibrils. GFP, green; phalloidin stained actin, red.

**Media File 6. Movie of a 3D reconstruction of Mhc^Wee-P26^-GFP integration into *bru1^M3^* mutant sarcomeres.**

Movie of an x,y,z-axis reconstruction from a confocal image of Mhc-weeP26-GFP in a *bru1^M3^* background. GFP signal is no longer restricted to the pre-myofibril myosin elements, and instead integrated across the sarcomere. GFP, green; phalloidin stained actin, red.

### Developmental upregulation of fibrillar genes is impaired in *bru1-IR* IFM

While investigating fiber-type specific exon misregulation, we noticed that genes and exons that are normally expressed preferentially in mature IFMs are downregulated in *bru1^M3^* IFMs (Fig S3 I). As CELF1 is known to promote embryonic splice events in vertebrate muscle (74), we wondered if developmentally-regulated transitions in gene expression and splicing are also disrupted in IFM lacking Bru1. To explore temporal-dependent dynamics in gene expression and exon use, we analyzed an mRNA-Seq time course in control and *bru1-IR* muscle at 24 h, 30 h and 72 h APF and 1 d adult. We confirmed via qPCR that *bru1-IR* (Mef2-Gal4 driven RNAi against *bru1*) results in a significant, 56-fold decrease in *bru1* mRNA levels in IFM (Fig S7 A). We noted that at all four timepoints, we could detect significant changes in gene expression and exon use in *bru1-IR* IFMs (Fig 4 A, Fig S4 A, Table S3). There was a dramatic increase in the number and magnitude of changes at both the gene and exon level in *bru1-IR* at 72 h APF and 1 d adult (Fig 4 A, Fig S4 A), which mirrors the increase in severity in cellular phenotypes in *bru1^M3^*IFMs from 80 h APF to 1 d adult (Fig 1 A, E, Fig 3 D, G). These results show that the number and severity of gene and exon regulatory defects progressively increases in *bru1-IR* IFM as development proceeds.

**Fig 4.**
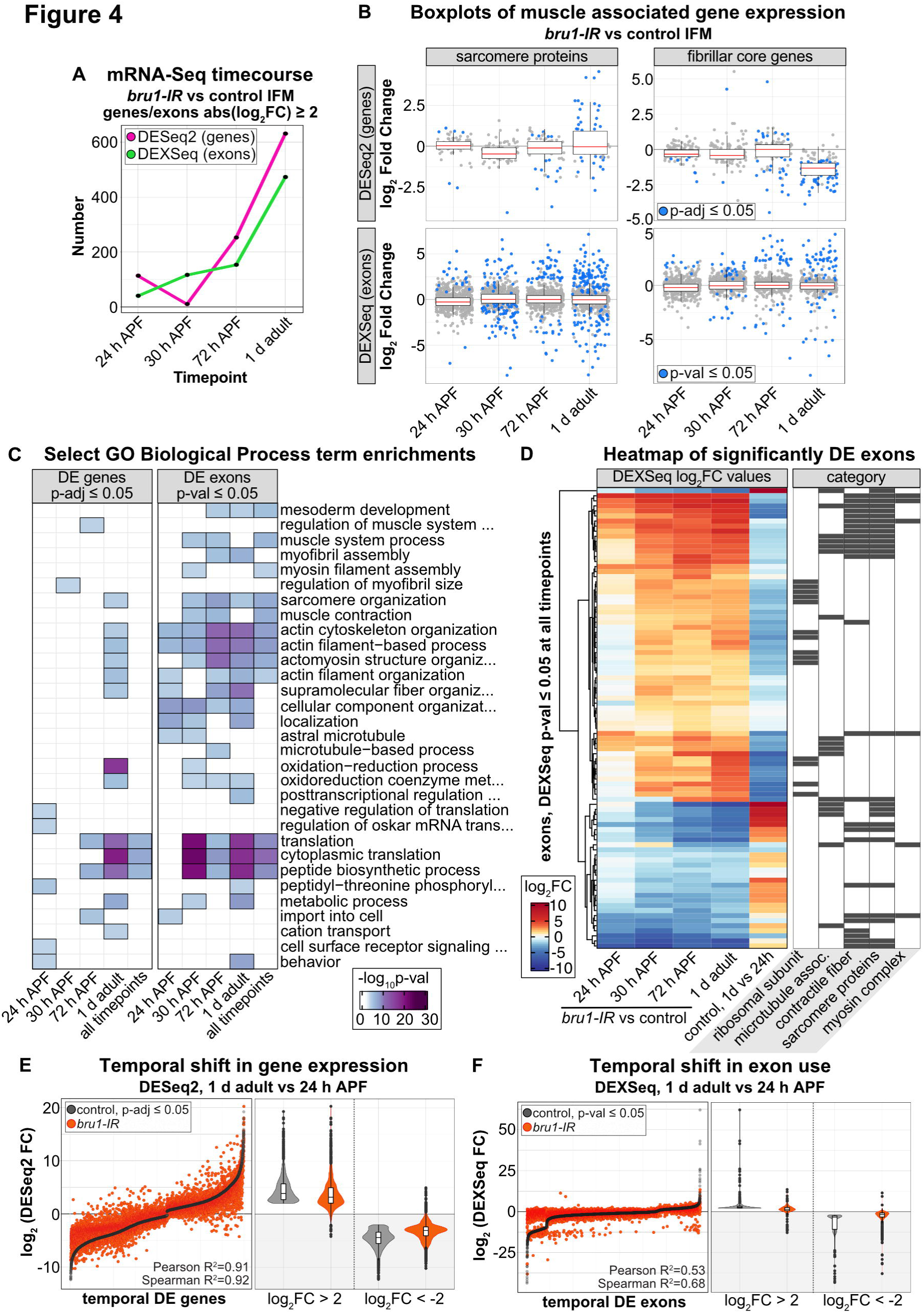
Knockdown of *bru1* disrupts temporal dynamics of gene expression and alternative splicing necessary for maturation of flight muscle. **(A)** Plot of the number of genes (magenta) and exons (green) significantly differentially expressed (p ≤ 0.05, abs(log_2_FC) ≥ 2) in *bru1-IR* versus control IFM at 24 h, 30 h, 72 h APF and in 1 d adult. **(B)** Boxplot of changes in gene expression (DESeq2) and exon use (DEXSeq) in sarcomere proteins and fibrillar muscle genes across the *bru1-IR* IFM time course. Blue dot denotes p ≤ 0.05. **(C)** Heatmap of select biological process GO term enrichments in significantly regulated genes (DESeq2, p-adj ≤ 0.05) and exons (DEXSeq, p-val ≤ 0.05) in *bru1-IR* IFM at 24 h, 30 h, 72 h APF or in 1 d adult, or at all four timepoints. **(D)** Heatmap (left) of all exons significantly DE (DEXSeq, p-val ≤ 0.05) at all timepoints in *bru1-IR* versus control IFM. The fifth column shows the temporal change in use of the same exons in *w^1118^* IFM from 24h APF to 1 d adult. Exons are identified as belonging to ribosomal subunit, microtubule associated, contractile fiber, sarcomere proteins or myosin complex gene categories (right, black boxes). **(E)** Left: Plot of the log_2_FC values of all genes differentially expressed (DESeq2, p-adj ≤ 0.05) in *w^1118^* IFM (black dots) from 24h APF to 1 d adult, ordered by control log_2_FC, and the corresponding change in the same genes in *bru1-IR* IFM (orange dots). Right: Violin plots comparing control (grey) and *bru1-IR* (orange) expression of strongly upregulated (log_2_FC ≥ 2) and downregulated (log_2_FC ≤ -2) temporal-switch genes. **(F)** Left: Plot of the log_2_FC value of all exons differentially expressed (DEXSeq, p-val ≤ 0.05) in *w^1118^*IFM (black dots) from 24h APF to 1 d adult, and the corresponding change in those same exons in *bru1-IR* IFM (orange dots). Right: Violin plots comparing exon use of strongly upregulated (log_2_FC ≥ 2) and downregulated (log_2_FC ≤ -2) temporal-switch exons in control (grey) and *bru1-IR* (orange) IFM.

*Drosophila* IFM has been shown to undergo a developmental transition in gene expression that is necessary to establish the stretch-activation mechanism characteristic of mature flight muscle (37). We next evaluated if this transition is disrupted in *bru1-IR* IFM. We evaluated specific subsets of genes previously shown to change across IFM development, including SPs, fibrillar muscle genes and actin cytoskeleton genes. While we saw downregulation of individual SPs and cytoskeleton genes, almost all core fibrillar muscle genes are strongly downregulated in 1 d adult *bru1-IR* IFM (Fig 4 B, Fig S4 B, C). This is also reflected in GO term enrichments across the time course. On the gene expression level, at pupal timepoints we found enrichment for terms such as “negative regulation of translation,” “cell surface receptor signaling pathway,” and “behavior” at 24 h APF, “regulation of myofibril size” at 30 h APF, and “regulation of muscle system development” and “translation” at 72 h APF (Fig 4 C). In 1 d adult bru1-IR IFM, we saw enrichment for cytoskeletal and muscle terms such as “sarcomere organization,” “actin cytoskeleton organization,” and “actomyosin structure organization,” as well as terms related to metabolism and translation such as “oxidation-reduction process,” “cytoplasmic translation” and “metabolic process.” To test genome-wide if this switch in gene expression during IFM maturation is disrupted, we identified all genes in our mRNA-Seq time course that are significantly regulated between 24 h APF and 1 d adult in control IFM, and plotted their temporal change in expression in *bru1-IR* IFM (Fig 4 E). This revealed that while *bru1-IR* IFM do undergo a temporal switch in gene expression, the switch is not as clean as in control IFM. The magnitude of up- or downregulation is often reduced in *bru1-IR* IFM, and a subset of genes show regulation in the opposite direction between 24 h APF to 1 d adult (Fig 4 E, Fig S4 A). We conclude that loss of Bru1 selectively impairs upregulation of fibrillar muscle genes but not the entire maturation program during IFM development.

### A developmental switch to mature splice isoforms is blocked in IFM lacking Bru1

Based on the developmental transition in gene expression, we hypothesized that a similar transition in alternative splicing exists in IFM. We next investigated if such a splicing transition exists, and if it is disrupted in IFM lacking Bru1. The Biological Process GO terms enriched in the hundreds of differentially expressed exons at all four timepoints in our mRNA-Seq time course (Fig 4 A) included “actin cytoskeleton organization,” “actin filament-based process”, “actomyosin structure organization,” “sarcomere organization” and “cytoplasmic translation” (Fig 4 C). We therefore started by looking at exon use dynamics in core fibrillar muscle genes and SPs, as several mature, IFM-specific isoforms of proteins in these categories have been reported (14,61,75,76). In total, at 24 h, 30 h or 72 h APF or at 1 d adult, we saw significant changes in 222 exons from 73 core fibrillar muscle genes and 413 exons from 56 SPs in *bru1-IR* IFM (Fig S4 D). Strikingly, exons that were upregulated in *bru1-IR* IFM were downregulated in control IFM between 24 h to 1 d adult, while exons that were downregulated in *bru1-IR* IFM were upregulated in control IFM between 24 h to 1 d adult.

To expand beyond SPs and fibrillar genes, we next identified a set of 91 exons that are significantly misregulated in at least three of four timepoints in *bru1-IR* IFM. These exons became more strongly misregulated as development proceeds and belonged to genes encoding ribosomal subunits, microtubule associated genes, SPs, contractile fiber and myosin complex genes (Fig 4 D). When we looked at the temporal change in use of these exons in control muscle from 24 h APF to 1 d adult, we found that all of the exons upregulated in *bru1-IR* IFMs are normally downregulated in 1 d adult muscle. Likewise, the exons downregulated in *bru1-IR* muscle are normally upregulated between 24 h APF and 1 d adult in control IFM (Fig 4 D). We then identified all exons in our mRNA-Seq time course that are significantly regulated between 24 h APF and 1 d adult in control IFM (Fig 4 F). We observed a clear temporal switch in exon use, reflecting mainly alternative splice events and alternative 3’ UTRs, but also alternative promoter use (Table S3). Strikingly, when we plotted the temporal change in use of these exons in *bru1-IR* IFM, we observed a near complete lack of developmental regulation. This analysis reveals that loss of Bru1 results in a block in the temporal shift in exon use during IFM development, and reveals that the underlying molecular defect in IFM lacking Bru1 is a failure to express mature, muscle-type specific splice isoforms.

To verify that changes observed in mRNA-Seq data reflect changes in protein expression, we analyzed the temporal dynamics of GFP-tagged reporters under control of endogenous regulatory elements for *wupA*, *Kettin*, *Clip190*, *Mlp60A* and *Mlp84B*. Although gene expression levels of *wupA*, *Kettin* and *Clip190* are consistent between control and *bru1-IR* across the mRNA timecourse (Fig S5 A), there are differences in the use of the isoform containing the exon where the GFP tag is inserted (Fig S5 B). *wupA-GFP*, which labels a termination used preferentially in tubular muscle (14) (Fig S2 D, I, J’, K’), is already visible in *bru1-IR* but not control IFM at 48 h and 72 h APF (Fig S5 C), confirming its missplicing throughout development in *bru1-IR* IFMs. Kettin encodes a short isoform of *sls* that is preferentially expressed in tubular muscle (77). Kettin-GFP is not expressed in control IFM, but is observed in *bru1-IR* IFM from 72 h APF (Fig S5 D). A GFP tag inserted in exon 29 of *Clip-190* is only expressed weakly in adult IFM in control, but is already expressed at 48 h and strongly at 72 h APF in *bru1-IR* IFM (Fig S5 E). Both Mlp60A and Mlp84B encode a single protein isoform that shows strong upregulation at the gene level in *bru1-IR* IFM from 72 h APF to 1 d adult. We can detect GFP expression of Mlp60A-GFP and Mlp84B-GFP in IFMs from 1 d adult *bru1-IR* flies, but not earlier timepoints at 48 h and 72 h APF (Fig S5 F, G). These data confirm that temporal regulatory dynamics observed at the mRNA level reflect temporal changes in protein and protein isoform expression in *bru1-IR* IFM. Taken together, our results significantly expand our understanding of the Bru1 phenotype and demonstrate that a block in developmental isoform expression, including increased expression of tubular-preferential isoforms, underlies the myofibril growth and hypercontraction defects observed during myofibril maturation in *bru1-IR* and *bru1^M3^* IFM.

### Loss of Bru1 leads to cytoskeletal organization defects at early stages of myogenesis

Although published Bru1 phenotypes in IFM are only reported after 48 h APF (14,36), our data above show that gene expression and splicing of structural genes are already abnormal in *bru1-IR* IFM from 24 h APF, prior to myofibril formation, and that abnormal sarcomere structure and contractility are already evident in *bru1^M3^* myofibers at 48 h APF. We therefore investigated the possibility that Bru1 also has a function in early IFM development. We extended our cross-section time course to include timepoints at 24 h, 26 h, 28 h, 30 h, 32 h and 34 h APF. Already at 24 h APF, we detected a defect in actin cytoskeleton organization in *bru1^M3^*myofibers (Fig 5 A). In wild-type *w^1118^* IFM at 24 h APF, F-actin is tightly organized at the sarcolemma, while in *bru1^M3^*myofibers it forms a more diffuse meshwork throughout the sarcoplasm that fails to condense as tightly as in wild-type (Fig 5 B, C). By 26 h APF in *w^1118^*myofibers, F-actin organizes into uniformly-distributed cables at the sarcolemma, which then subdivide and migrate towards the center of the myofiber from 28-30 h APF. In *bru1^M3^* this process is abnormal, with F-actin organizing into many smaller, unequally distributed cables with impaired migration (Fig 5 C, D). Both *w^1118^* and *bru1^M3^* myofibers undergo myofibrillogenesis from 30-34 h APF, but there are fewer myofibrils present in *bru1^M3^* myofibers by 34 h APF and those myofibrils are significantly larger in diameter than *w^1118^*myofibrils (Fig 5 C, E, F). Progressive organization of the actin cytoskeleton during IFM myogenesis leads to myofiber compaction in addition to generating the tension necessary to drive myofibrillogenesis (58,70,71). At both 24 h and 30 h APF, myofibers are longer in *bru1^M3^* than in *w^1118^* (Fig 5 G, H), demonstrating that IFM compaction is mildly impaired and consistent with an actin cytoskeleton defect in myofibers lacking Bru1. We conclude that Bru1 is necessary during early IFM development to regulate cytoskeletal organization dynamics that support proper myofibril formation.

**Fig 5.**
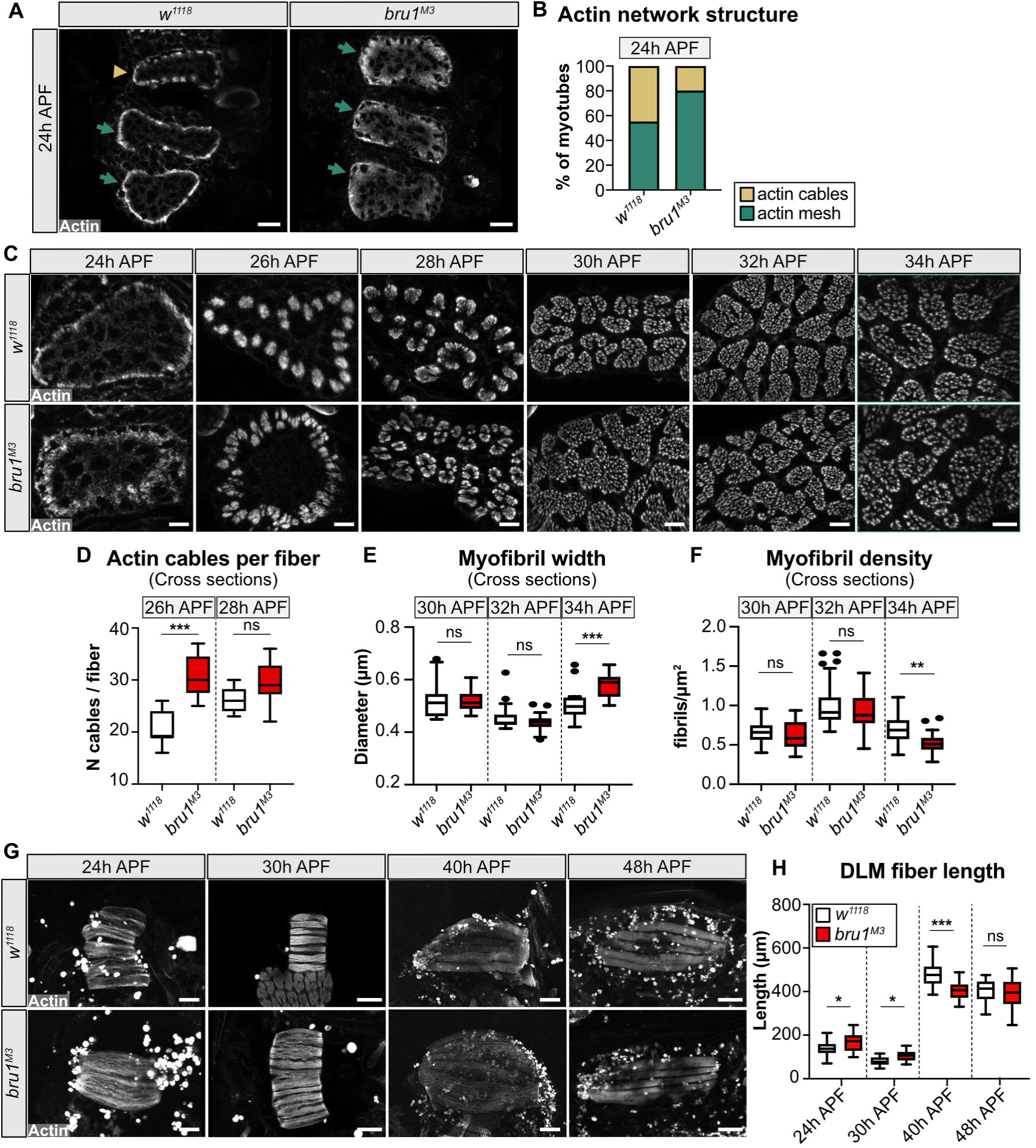
Early cytoskeletal rearrangement and myofiber compaction are abnormal in *bru1* mutant IFM. **(A)** Confocal images of DLM cross-sections at 24 h APF. Phalloidin stained F-actin (grey) reveals progressive condensation of the actin network into cables (arrow head) near the sarcolemma in *w^1118^*, but a meshwork (arrow) in *bru1^M3^*. Scale bar = 10 μm. **(B)** Quantification of actin network structure in (A). N > 10 for each genotype. **(C)** Cross-section time-course of early cytoskeletal rearrangements and pre-myofibril formation in *w^1118^* and *bru1^M3^* at 24 h, 26 h, 28 h, 30 h, 32 h, 34 h APF. Scale bar = 5 μm. Irregular actin condensation at 24 h and cable splitting at 26 h and 28 h APF is evident in *bru1^M3^*, prior to pre-myofibril formation at 30-32 h. **(D)** Quantification of actin cable number per myotube in *w^1118^*and *bru1^M3^* at 26 h and 28 h APF. Significance determined by ANOVA and post-hoc Tukey (ns, not significant; ***, p-val < 0.001). **(E-F)** Quantification of myofibril width (E) and density (F) in *w^1118^* and *bru1^M3^*at 30 h, 32 h and 34 h APF. The first sarcomere growth defects in *bru1^M3^* are already detected at 34 h APF, when the pre-myofibrils are fully formed. Significance determined by ANOVA and post-hoc Tukey (ns, not significant; **, p-val < 0.01; ***, p-val < 0.001). All boxplots are shown with Tukey whiskers, outlier data points marked as black dots. **(G)** Confocal Z-stack images of DLMs in *w^1118^* and *bru1^M3^* at 24 h, 30 h, 40 h and 48 h APF. DLM fibers undergo compaction at 30 h APF, followed by re-extension and fiber growth. Phalloidin stained actin, grey; Scale bar at 24, 30 h APF = 50 μm, at 40, 48 h APF = 100 μm. **(H)** Quantification of the DLM fiber length in (G). *bru1^M3^* DLM fibers fail to fully compact. Significance determined by ANOVA and post-hoc Tukey (ns, not significant; *, p-val < 0.05; ***, p-val < 0.001).

### Temporal restricted RNAi reveals a requirement for Bruno1 in early and late myogensis

The data we present above suggest that Bru1 is required both during early and late stages of IFM myogenesis. To test the temporal requirement of Bru1 in muscle development, we performed RNAi with a panel of five Gal4 enhancer lines with distinct temporal expression patterns in IFM (Fig S6 A). In contrast to Mef2-Gal4, which is expressed in all muscle throughout development, salm-Gal4, Act88F-Gal4, UH3-Gal4 and Fln-Gal4 are expressed in IFM from approximately 16 h APF, 24 h APF, 34 h APF and 56 h APF, respectively (76,78–80). Him-Gal4 is expressed in myoblasts and is downregulated in IFM by 30 h APF (37). *bru1-IR* led to a strong reduction in *bru1* mRNA with all drivers tested (Fig S7 A), but did not impair adult viability or eclosion (Fig S6 B). We then compared phenotypes of *bru1-IR* driven by Gal4 lines active during early myogenesis (*bru1-IR^Him^*, *bru1-IR^salm^*, *bru1-IR^Act88F^*) to those active after myofibrillogenesis is completed (*bru1-IR^UH3^* and *bru1-IR^Fln^*).

Knockdown of *bru1* during early myogenesis with Him-Gal4, salm-Gal4 or Act88F-Gal4 resulted in severe behavioral and structural defects. With all three drivers, flies were flightless (Fig 6 B, Fig S6 C) and the majority of IFMs were ripped or completely detached (Fig 6 B-D, Fig S6 C, D). The percent of detached myofibers between 90 h APF and 1 d adult increased from 39.5% to 72.5%, 16.3% to 51.4% and 37.5% to 83.3% in *bru1-IR^Him^*, *bru1-IR^salm^* and *bru1-IR^Act88F^*, respectively, reflecting the progressive hypercontraction phenotype (Fig 6 C, D, Fig S6 D, E-N’). *bru1-IR^Him^*, *bru1-IR^salm^* and *bru1-IR^Act88F^*also produced similar myofibril and sarcomere phenotypes characterized by short, thick sarcomeres and torn myofibrils (Fig 6 D, Fig S6 E-N’). At both 90 h APF and 1 d adult, *bru1-IR* sarcomeres are significantly shorter and wider than control sarcomeres (Fig 6 E, F, Fig S6 F, G). Interestingly, although Him-Gal4 turns off at 30 h APF, while salm-Gal4 and Act88F-Gal4 are expressed continuously starting from 16 h and 24 h APF, respectively, they all produce a similar *bru1-IR* phenotype. We confirmed that *bru1-IR^Him^* results in a loss of Bru1 protein at 24 h and 30 h APF, but from 48 h APF Bru1 protein is present in the nuclei of *bru1-IR^Him^* flies (Fig S7 M-Q). This data shows that decreased expression of Bru1 during early stages of IFM development is sufficient to produce severe myofibril and sarcomere phenotypes.

**Fig 6.**
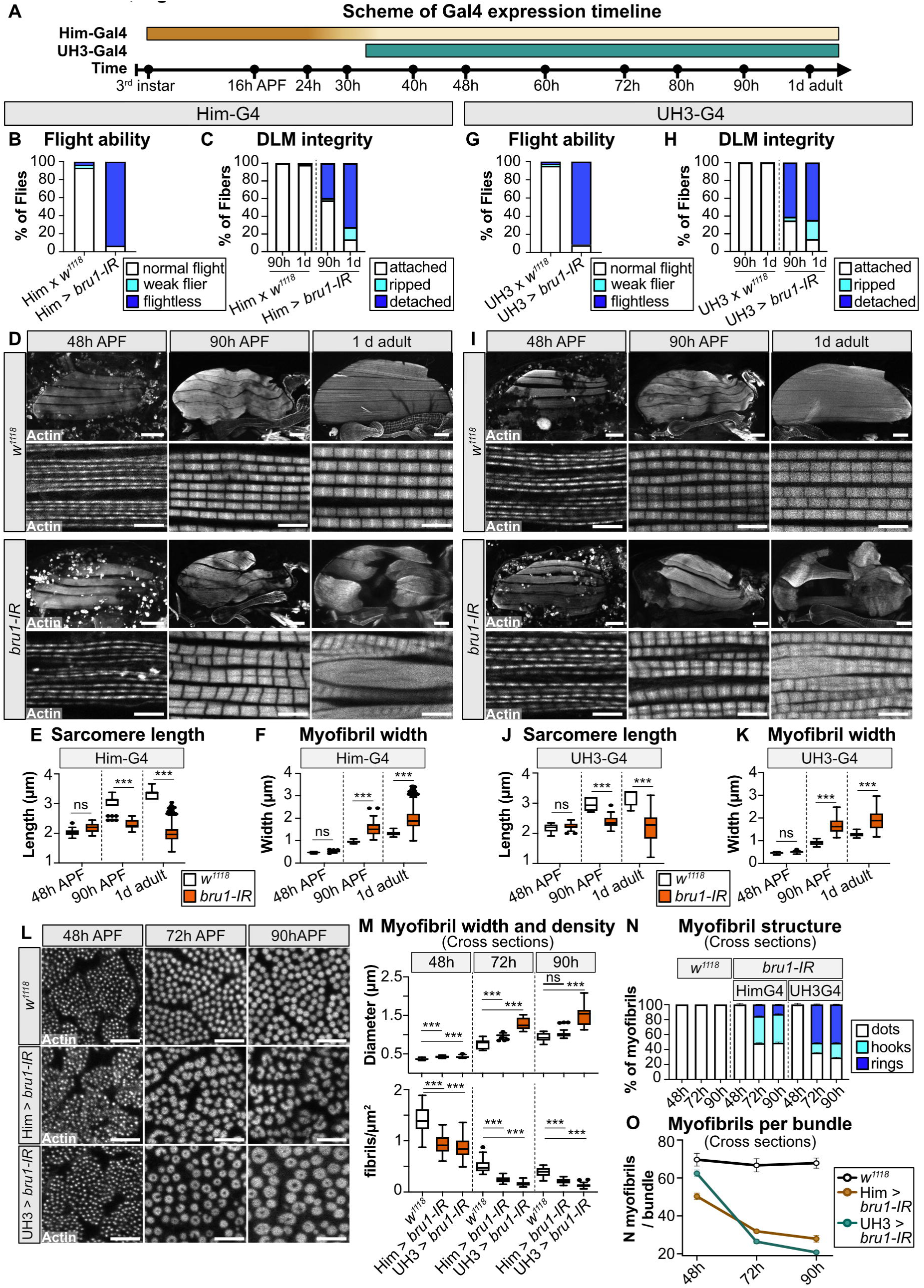
Temporal-restricted RNAi demonstrates a functional requirement for Bru1 during early myogenesis. **(A)** Scheme of Him-Gal4 and UH3-Gal4 expression timing during IFM myogenesis. Gradient colour of the bar indicates the strength of expression. Time-points as marked. **(B)** Quantification of flight ability in *Him*-Gal4 driven *bru1-IR* (*bru1-IR^Him^*) (N > 130). **(C)** Quantification of myofiber ripping and detachment phenotypes in control and *bru1-IR^Him^*at 90 h APF and 1d adult (N > 40). **(D)** Confocal projections of control and *bru1-IR^Him^* hemithoraxes (upper, scale bar = 100 μm) and single-plane images of myofibrils (lower, scale bar = 5 μm) at 48 h and 90 h APF and 1 d adult. **(E-F)** Quantification of sarcomere length (E) and myofibril width (F) from (D). Boxplots are shown with Tukey whiskers, outlier data points marked as black dots. Significance determined by ANOVA and post-hoc Tukey (ns, not significant; ***, p-val < 0.001). **(G)** Quantification of flight ability in *bru1-IR^UH3^* (N > 110). **(H)** Quantification of myofiber integrity in control and *bru1-IR^UH3^*at 90 h APF and 1d adult (N > 40). **(I)** Confocal projections of hemithoraxes (upper, scale bar = 100 μm) and single-plane images of myofibrils (lower, scale bar = 5 μm) in control (top) and *bru1-IR^UH3^* (bottom) at 48 h and 90 h APF and 1 d adult. The severity of the *bru1*-RNAi associated myofibril phenotype is comparable between (D) and (I). **(J-K)** Quantification of sarcomere length (J) and myofibril width (K) from (I). Significance determined by ANOVA and post-hoc Tukey (ns, not significant; ***, p-val < 0.001). **(L)** Single-plane cross-section images of DLM myofibrils in control *w^1118^*, *bru1-IR^Him^*and *bru1-IR^UH3^* at 48 h, 72 h and 90 h APF. Scale bar = 5 μm. **(M)** Quantification of myofibril width (top) and density (bottom) in (L). Significance determined by ANOVA and post-hoc Tukey (ns, not significant; ***, p-val < 0.001). **(N)** Quantification of myofibril structure from (L), showing the ratio of normal dot morphology (white) to abnormal hook (cyan) and ring (blue) structures. Fewer hooks and rings form in early expressed *bru1-IR^Him^* as compared to *bru1-IR^UH3^*. Error bars = SEM. **(O)** Quantification of mean myofibril number per bundle in (L). *bru1-IR^UH3^* IFM forms the correct number of myofibrils while *bru1-IR^Him^* IFM does not, but myofibril fusion is more extensive after 48 h APF in *bru1-IR^UH3^*. Error bars = SEM. F-actin in (D, I, L) stained with phalloidin.

We then examined *bru1* knockdown phenotypes in the UH3-Gal4 and Fln-Gal4 drivers that are expressed after myofibrillogenesis. While *bru1-IR^UH3^* produces flightless flies with ripped and detached IFM myofibers (Fig 6 G, H, I), *bru1-IR^Fln^*flies are able to fly and have intact IFMs (Fig S6 C, D, E). Although *bru1* mRNA levels are significantly decreased by both *bru1-IR^UH3^* and *bru1-IR^Fln^* (Fig S7 A), Bru1 protein still detected in *bru1-IR^Fln^* myofibers at 90 h APF, but cannot be detected in *bru1-IR^UH3^* myofibers (Fig S7 D-H). We therefore focused our subsequent analysis on *bru1-IR^UH3^*. Like *bru1-IR^Him^*, *bru1-IR^salm^* and *bru1-IR^Act88F^*, *bru1-IR^UH3^* produces sarcomeres that are shorter and thicker than control sarcomeres at 90 h APF and 1 d adult, as well as torn myofibrils and progressive loss of sarcomere architecture in 1 d adults (Fig 6 I, J, K). However, when examined in cross-section, *bru1-IR^UH3^* produces stronger radial growth defects than *bru1-IR^Him^* (Fig 6 L). *bru1-IR^UH3^*myofibrils are significantly wider in diameter than control or *bru1-IR^Him^* myofibrils at both 72 h and 90 h APF (Fig 6 M), and proportionally more myofibrils appear as rings instead of hooks or dots already at 72 h APF (Fig 6 N). In addition, although *bru1-IR^UH3^* IFMs have a near wild-type number of 62 ± 12 myofibrils per bundle at 48 h APF, by 72 h APF the number of 26 ± 6 myofibrils per bundle is less than the 32 ± 7 observed with *bru1-IR^Him^* (Fig 6 O). This suggests that the recovering level of Bru1 protein after 48 h APF in *bru1-IR^Him^* promotes a somewhat less severe radial overgrowth phenotype than the absence of Bru1 protein from 48 h APF onwards in *bru1-IR^UH3^*. These data support a requirement for Bru1 during myofibril maturation after 48 h APF to regulate sarcomere growth dynamics.

### Levels of Bru1 expression during early myogenesis impact myofibril formation and growth

Our data indicate that phenotypes related to Bru1 function in early and late myogenesis may be separable, where early function defines the number and organization of myofibrils while later function regulates sarcomere radial and lateral growth. To test this hypothesis, we performed temporally-regulated rescue experiments. We generated a UAS-Bru1 rescue construct (Fig S8 A), and then optimized expression conditions with our Gal4 panel, as overexpression with Mef2-Gal4 causes severe phenotypes (14,36). Expression of UAS-Bru1 with salm-Gal4 lead to embryonic lethality (Fig S8 B). Overexpression with Act88F-Gal4 caused loss of flight ability, severely degraded myofibrils and loss of IFM myofibers (Fig S8 C-F). UH3-Gal4 mediated overexpression of Bru1 caused loss of flight ability due to abnormally short, thin and trapezoidal-shaped sarcomeres (Fig S8 C-G). Overexpression with the two Gal4 drivers with the narrowest expression windows, Him-Gal4 and Fln-Gal4 (Fig S6 A), produced surviving flies (Fig S8 A) with intact IFMs and enabled us to evaluate Bru1 function in the early 0-30 h APF and late 56 h APF to adult time windows.

To evaluate Bru1 function during early IFM development, we performed overexpression and rescue experiments with Him-Gal4. We first confirmed that in a *bru1^M3^* background, Him-Gal4 drives expression of Bru1 at 24 h APF, but not at 48 h or 90 h APF (Fig S9 A, B), demonstrating the temporal-restriction of Bru1 expression to early myogenesis. Overexpression of Bru1 in a wild-type background is sufficient to produce short, thick sarcomeres and loss of flight ability, but IFM myofibers remain attached (Fig 7 A-E, Fig S9 C-F). This demonstrates that overexpression of Bru1 produces the same sarcomere phenotype as loss-of-function, indicating that Bru1 is dosage sensitive. In cross-sections, overexpression of Bru1 with Him-Gal4 leads to a reduced myofibril density, a decreased number of myofibrils per bundle and myofibrils of variable thickness at 72h and 90 h APF, but is not sufficient to produce hollow myofibrils (Fig 7 F-I, Fig S9 I). Unexpectedly, IFMs overexpressing Bru1 have a prominent hole in the middle of the myofiber at 48 h APF, which we interpret to reflect failed migration of actin cables to the center of the fiber from 28-30 h APF (Fig 5 C). These data demonstrate that the dosage of Bru1 before 30 h APF is crucial to regulate the number and regularity of myofibrils, and can further influence sarcomere growth during later developmental stages.

**Fig 7.**
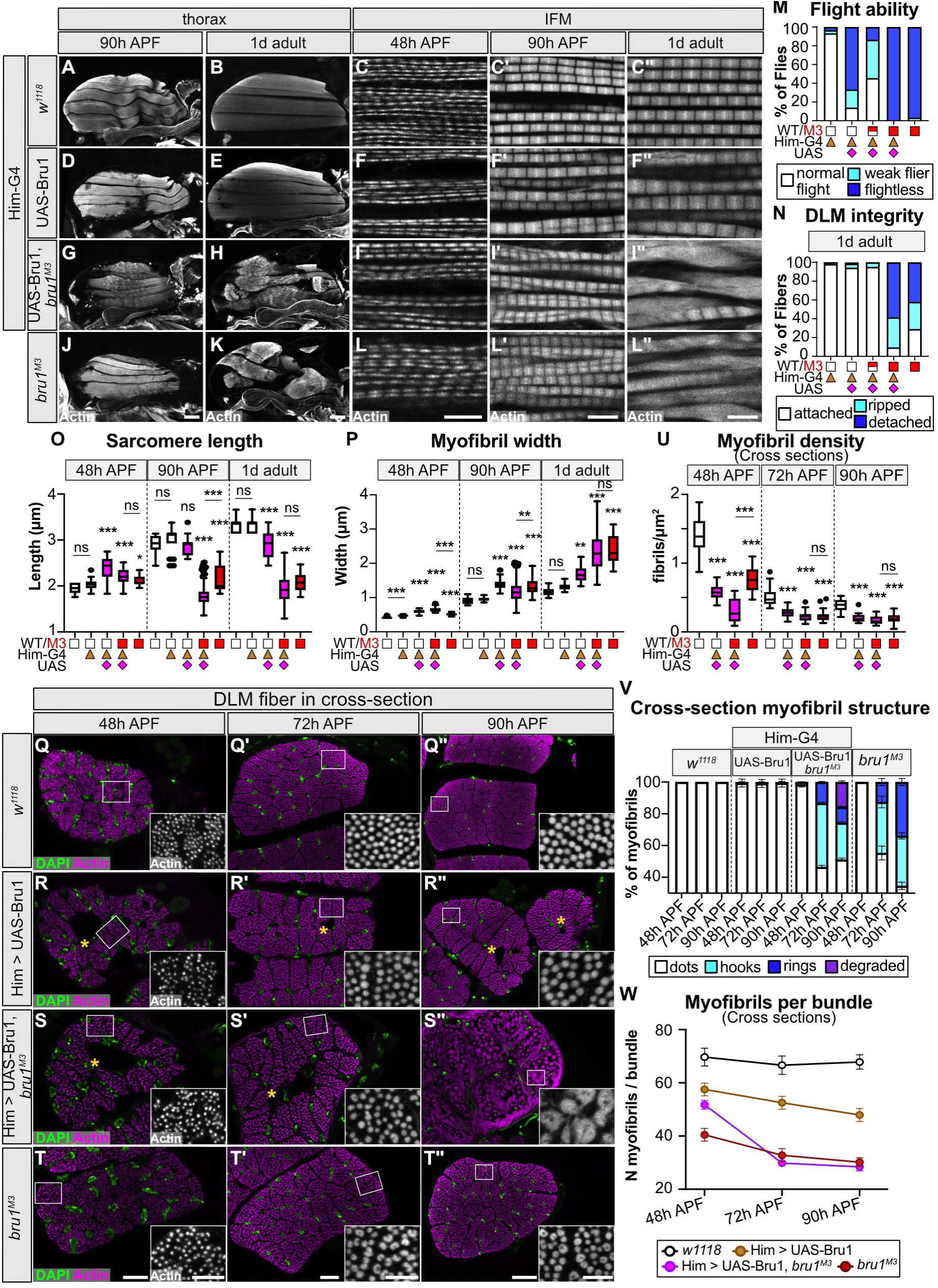
Expression of Bru1 restricted to early myogenesis fails to rescue and exacerbates *bru1^M3^* myofibril phenotypes. **(A-L”)** Confocal projections of hemithorax (scale bar = 100 μm) and single-plane images of myofibril structure (scale bar = 5 μm) in control (A-C”); Bru1 overexpression (*Him*-Gal4 > UAS-Bru1, D-F”); early temporal rescue (*Him*-Gal4 > UAS-Bru1, *bru1^M3/M3^*, G-I”); and mutant *bru1^M3^*(J-L”). Time points at 48 h, 90 h and 1 d adult as labelled. Phalloidin stained actin, grey. **(M-N)** Quantification of flight ability (M) and DLM fiber integrity (N) in (B, E, H, K). Genotypes labeled at the bottom: wildtype, white square; *bru1^M3^*, red square; *Him*-Gal4, orange triangle; UAS-Bru1, magenta diamond. N > 40 fibers for each genotype. **(O-P)** Quantification of sarcomere length (O) and myofibril width (P) in (C, F, I, L). Boxplots are shown with Tukey whiskers, outlier data points marked as black dots. Significance determined by ANOVA and post-hoc Tukey (ns, not significant; *, p-val < 0.05; **, p-val < 0.01; ***, p-val < 0.001). **(Q-T”)** Cross-section image of myofibril structure at 48 h, 72 h and 90 h APF in control (Q), Bru1 overexpression (R), early temporal rescue (S), and mutant *bru1^M3^* IFM (T). Magnified image of selected area (white rectangle) shown in lower right corner. Overexpression of Bru1 with *Him*-Gal4 produces holes in the center of IFM myofibers (yellow asterisks) and cannot rescue later formation of hollow myofibrils; DAPI, green; phalloidin stained actin, magenta; Scale bar = 10 μm (Q-T, Q’-T’), 20 μm (Q”-T”) and 5 μm (zoom-in sections). **(U-V)** Quantification of myofibril density (U) and myofibril structure (V) in (Q-T”). Genotypes, boxplot and significance as above. Error bars in V = SEM. **(W)** Quantification of mean myofibril number per fiber bundle in (Q-T”). *Him*-Gal4 driven Bru1 can partially rescue the number of myofibrils formed in *bru1^M3^* myofibers before 48 h APF, but cannot rescue myofibril fusion and hollow myofibril formation after 48 h APF. Error bars = SEM.

We then investigated if Him-Gal4 mediated expression of Bru1 is sufficient to rescue the *bru1^M3^* phenotype. Instead of rescuing, expression of Bru1 during early developmental stages actually increases the severity of the *bru1^M3^* phenotype. Rescue flies are flightless, and show greater proportions of flies with ripped or detached IFMs at both 90 h APF and 1 d adult (Fig 7 A-C, Fig S9 C-D). Sarcomeres in rescue IFMs are significantly shorter than both control and *bru1^M3^*sarcomeres at 90 h APF (Fig 7 D), are as wide as mutant sarcomeres (Fig 7 E), and show myofibril tearing and loss of sarcomere architecture in 1 d adults (Fig 7 A). In cross-section, Him-Gal4 > Bru1 rescue flies form hollow myofibril structures that are larger and more irregular than those formed by *bru1^M3^* myofibrils at 90 h APF (Fig 7 F). Interestingly, while the myofibril density and developmental progression of hollow myofibril formation mirrors that observed in *bru1^M3^* (Fig 7 G, H), rescue flies show a significant reduction in the number of myofibrils per bundle from 48 h to 72 h APF (Fig 7 I), indicating that Bru1 expression may partially rescue some early myofibril defects. Rescue flies also have a hole in the middle of the IFM myofiber that is evident at 48 h, 72 h and 90 h APF (Fig 7 F). We confirmed the presence of this hole using hematoxylin and eosin staining, as well as the absence of such a hole in wild-type or *bru1^M3^* myofibers (Fig S9 G-H). Taken together, these data demonstrate: 1) that Bru1 has a function during early stages of IFM myogenesis; 2) myofibril phenotypes are sensitive to Bru1 expression level before 48 h APF; 3) loss and gain of Bru1 can produce similar phenotypes and 4) misregulation of Bru1 in early development also impacts later stages of myofibril maturation.

### Rescue of Bru1 during late development is insufficient to rescue myofibril structural defects

To evaluate Bru1 function during late IFM development, we performed overexpression and rescue experiments with Fln-Gal4. We first verified by antibody staining that at 90 h APF in a *bru1^M3^*background, Him-Gal4 drives expression of UAS-Bru1 (Fig S10 A, B). Overexpression of Bru1 with Fln-Gal4 in a wild-type background was sufficient to produce flightless flies, but myofibers remained attached and did not display hypercontraction-related tearing (Fig 8 A-C, Fig S10 C). Although myofibril and sarcomere structure was largely intact, sarcomeres were significantly shorter and thinner than in the control and 52.4% were frayed in 1 d adults (Fig 8 A, D-F, Fig S10 C-E). We conclude that overexpression of Bru1 during late IFM development with Fln-Gal4 is sufficient to produce flightlessness due to mild sarcomere defects, but not strong enough to reproduce the *bru1^M3^* phenotype.

**Fig 8.**
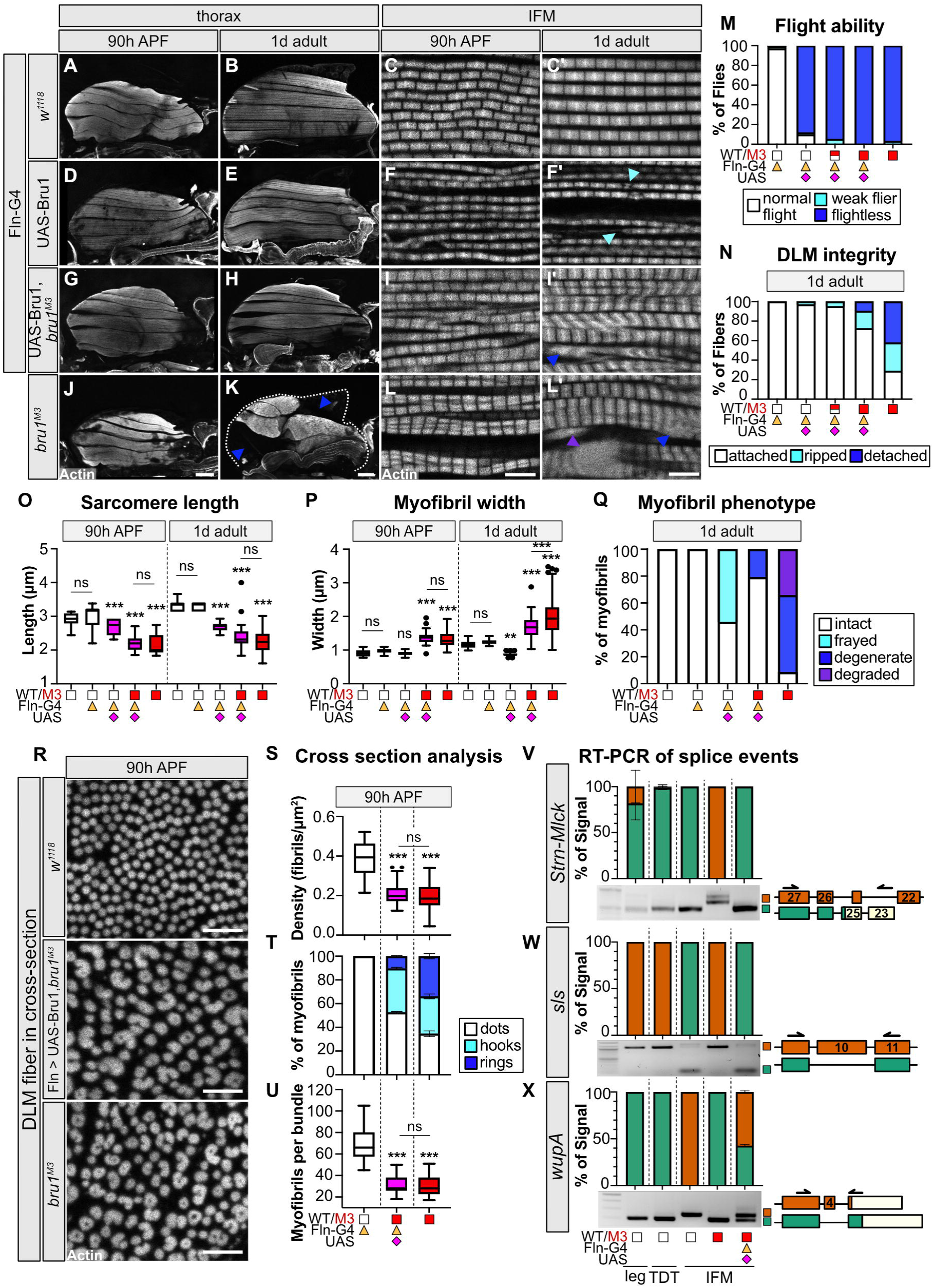
Expression of Bru1 restricted to late myogenesis partially rescues *bru1^M3^* myofibril phenotypes and restores alternative splicing defects. **(A-L’)** Confocal projections of hemithorax and single-plane images of myofibril structure at 90 h APF and 1 d adult in control (A-C’); Bru1 overexpression (*Fln*-Gal4 > UAS-Bru1, D-F’); late temporal rescue (*Fln*-Gal4 > UAS-Bru1, *bru1^M3/M3^*, G-I’); and mutant *bru1^M3^* (J-L’). The severity of myofiber detachment and torn myofibril phenotypes is partially rescued in *Fln*-Gal4 > UAS-Bru1, *bru1^M3^* IFM. Dashed line outlines the thorax boundary in (K). Frayed (cyan), degenerate (blue), or degraded (purple) myofibrils, arrowheads; Phalloidin stained actin, grey; Scale bar = 100 μm (thorax); scale bar = 5 μm (IFM). **(M-N)** Quantification of flight ability (M) and DLM fiber integrity (N) in (B, E, H, K) (N > 50). Genotypes labeled at the bottom: wildtype, white square; *bru1^M3^*, red square; *Fln*-Gal4, orange triangle; UAS-Bru1, magenta diamond. **(O-P)** Quantification of sarcomere length (O) and myofibril width (P) in (C-L’). Boxplots are shown with Tukey whiskers, outlier data points marked as black dots. Significance determined by ANOVA and post-hoc Tukey (ns, not significant; **, p-val < 0.01; ***, p-val < 0.001). **(Q)** Quantification of myofibril phenotypes present in (C’-L’) (N > 70). **(R)** Cross-section images of myofibril structure in DLM from control, late temporal rescue, and mutant *bru1^M3^*at 90 h APF. Phalloidin stained actin, grey; Scale bar = 5 μm. **(S-U)** Quantification of cross-section myofibril density (S), myofibril structure (T), and number of myofibrils per bundle (U) in (R). Fewer hollow myofibrils (rings) develop in *Fln*-Gal4 > UAS-Bru1, *bru1^M3^*IFM by 90 h APF. Significance determined as above, error bars in (T) = SEM. **(V-X)** RT-PCR verification of Bru1 regulated alternative splice events in *Strn-Mlck* (V), *sls* (W) and *wupA* (X). Representative gel images and quantification of percent exon use in control fibrillar IFM, tubular leg and jump (tergal depressor of the trochanter, TDT) muscle, and mutant *bru1^M3^* or late rescue *Fln*-Gal4 > UAS-Bru1, *bru1^M3^* IFM. Error bars = SD. Scheme on the right of alternative isoforms with primer locations, color coding consistent between scheme and bar plot. 3’-UTR regions in light beige. Exon numbering according to FB2021_05.

We then evaluated if expression of Bru1 from 56 h APF with Fln-Gal4 can rescue the *bru1^M3^* phenotype. Strikingly, whereas 71.0% of myofibers are ripped or detached in 1 d adult *bru1^M3^* flies, only 27.4% are ripped or detached in Fln-Gal4 rescue flies (Fig 8 A, C). In addition, while 91.5% of myofibrils are degraded, degenerate or frayed in *bru1^M3^* IFM, only 20.7% of myofibrils are affected in the rescue (Fig 8 F). However, the rescue flies are still flightless and their sarcomeres are also significantly shorter and thicker than control sarcomeres (Fig 8 A, B, D, E). Cross-section analysis revealed that hollow myofibril formation in the Fln rescue flies proceeds to the same extent as in *bru1^M3^* IFM, and myofibril density, the percent of myofibrils converted to hook and ring-like structures, as well as the number of myofibrils per bundle is not significantly different from the *bru1^M3^* mutant (Fig 8 G-J). This indicates that expression of Bru1 with Fln-Gal4 is sufficient to produce a partial rescue of hypercontraction-related myofiber tearing and loss of sarcomere architecture, but is insufficient to rescue sarcomere and myofibril growth defects. To further test this conclusion, we performed semi-quantitative RT-PCR to validate the rescue of individual alternative splice events we have previously shown to be regulated by Bru1 (14,38). We found that events in *Strn-Mlck*, *sls*, *Tm1* and *Mhc* are completely rescued (Fig 8 K, L, Fig S10 I, J), while events in *wupA* and *Zasp52* are partially rescued (Fig 8 M, Fig S10 H). We conclude that expression of Bru1 with Fln-Gal4 is sufficient to restore Bru1 mediated splicing of key structural genes and partially alleviate hypercontraction-related phenotypes in IFM. However, this late-stage rescue cannot repair pre-existing cytoskeletal structural and growth defects leading to a continued imbalance in growth in sarcomere length and width, abnormal radial growth of the myofibril and associated functional deficits.

## Discussion

CELF proteins are important RNA regulators in muscle of both vertebrates and insects. Here, employing the power of the *Drosophila* model system, we have dissected the pleiotropic Bru1 phenotype in IFM and propose the following developmental model explaining how misregulation of CELF proteins leads to muscle structural defects. In the developing myotube and nascent myofiber, Bru1 is required to promote actin cytoskeletal rearrangements that influence the number, organization and size of myofibrils that will be formed (Figs 1, 3, 5). Bru1 regulates isoform expression patterns that influence myofiber compaction dynamics during myofibrillogenesis, as well as the frequency and strength of spontaneous contractions after myofibril formation (Figs 2, 4, 5). As development proceeds, Bru1 enables a genome-wide switch to production of mature isoforms in structural and cytoskeletal regulatory genes (Fig 4). In *bru1* mutants, disruption of this switch leads to an imbalance in sarcomere growth in length and width and a pronounced misregulation of radial growth that alters the dynamics of actin and myosin incorporation into sarcomeres and drives myofibril fusion and the formation of hollow myofibrils (Fig 3, 6, 7). Further, loss of IFM-specific isoforms produces abnormal actomyosin contractility throughout myofiber development and a terminal hypercontraction phenotype that results in adult myofiber loss (Fig 1). Due to this developmental progression, rescue of Bru1 during late development can restore alternative splice events and partially alleviate hypercontraction and myofiber loss, but cannot correct pre-existing structural defects to restore sarcomere structure or muscle function (Fig 7, 8). Our model provides novel insight into the multiple stages of myofibrillogenesis regulated by Bru1 and the breadth of the fiber-type differentiation processes regulated by CELF proteins.

Developmentally regulated exons in individual sarcomeric genes such as vertebrate cardiac Troponin T (cTnT) (81,82), *Drosophila* Myosin heavy chain (Mhc) (83), and Titin-like proteins (84,85) are well characterized. More recently, RNA-sequencing technologies have revealed temporal shifts from embryonic to post-natal splice isoform expression in hundreds of cytoskeletal, calcium-regulatory and signaling genes in vertebrate heart and skeletal muscles (21,27,86). Although a recent study demonstrated that *Drosophila* IFMs undergo a transcriptional switch to enable myofiber maturation (37), genome-wide evidence for a similar switch in alternative splicing has been lacking. Here we identify a genome-wide developmental switch in exon use and gene isoform expression in IFMs (Fig 4). This switch is extensive and conservatively involves around 3,000 exons from 2,000 genes (Table S3), including cytoskeletal and sarcomeric proteins, metabolic genes and the translation machinery. These enrichment categories are consistent with similar changes in cytoskeletal, signaling and calcium regulatory proteins during maturation of vertebrate muscle (21,86,87). This indicates that sarcomere assembly and maturation involves distinct phases with specific physiological requirements, consistent with existing models of myofibrillogenesis (9,37,40), and global changes in isoform expression enable muscles in both vertebrates and insects to proceed through developmental transitions. The regulation of alternative splicing and more generally RNA processing is thus a conserved mechanism to fine-tune cytoskeletal dynamics, cellular metabolism, intracellular transport and protein expression.

While CELF1 and CELF2 are required during early mouse and chick myogenesis to promote embryonic splicing patterns in heart and skeletal muscle (18,19,21,88), previous studies on Bru1 in *Drosophila* IFMs only reported phenotypes after 48 h APF during later phases of myofibril maturation (14,36). Here we showed that Bru1 in flies also has a function in early IFM myogenesis before 48 h APF to regulate cytoskeletal rearrangement and myosin contractility (Figs 4-8), demonstrating a conserved requirement for CELF family function during early myogenesis. Our data show that like other CELF proteins, Bru1 regulates alternative splicing of hundreds of exons (Fig 4). In addition to a nuclear role in alternative splicing, CELF proteins have been shown to have cytoplasmic roles in regulating mRNA decay and translation (18,63,89,90). Based on the low or anti-correlation between mRNA-Seq and whole proteome mass spectrometry data for many peptides, our data also suggests a possible role for Bru1 in regulating mRNA stability or translation in IFM (Fig 1, Fig S1). Further studies are required to determine if this effect is direct, driven by changes in alternative exon or 3’-UTR use in target genes or indirect through interaction with other RBPs such as Rbfox1 (38). In vertebrates, downregulation of CELF1 and concurrent upregulation of MBNL1/2 promotes a transition to mature patterns of alternative splicing (21,23,24,91). It is still unclear if the same temporal regulatory relationship exists between Bru1 and Muscleblind (mbl) in *Drosophila*. Mbl is expressed in both larval and adult muscle, is necessary for z-disc and myotendinous junction maturation in embryonic muscle, and regulates both sarcomere growth and myosin contractility (13,92–94), but the temporal requirement for Mbl function as well as Mbl target genes remain to be determined. Our discovery of an early requirement for Bru1 in IFM development strengthens the use of *Drosophila* as a model to examine the molecular mechanism of CELF function as well as co-regulatory interactions with other RBPs.

The function of Bru1 before myofibrillogenesis as well as during stages of sarcomere growth and myofibril maturation (Figs 1-6) demonstrates a requirement for Bru1 throughout IFM development. This is consistent with previous work showing that Bru1 is expressed selectively and continuously in the IFMs and promotes fiber-type specific splice events in structural genes necessary to establish adult IFM contractile properties (14,36,38). Muscle-specific splicing factors are also found in vertebrates, for example RBM24 (95,96) or the Fragile-X like protein isoform FXR1P82,84 (97), and enhance splicing or stability of select isoforms of cytoskeletal genes including *MyoG*, *cTnT, eMHC, Mef2d and Naca* to promote muscle differentiation (96,98,99). CELF1 is also thought to play a role in fiber-type differentiation, as overexpression of either CELF1 or a dominant-negative CELF protein is sufficient to change the ratio of slow to fast fiber types (29,100). Beyond CELF1, expression of other CELF proteins including CELF2 (ETR-3) and CELF4 have been shown to increase as mouse thigh and heart muscle develop (23,27). These CELF proteins also regulate alternative splicing of sarcomere gene exons (27,101–103), for example CELF4 promotes splicing of mature isoforms of chicken β-tropomyosin (104). CELF2 expression in chicken and mouse shifts from preferential production of an embryonic 52 kDa isoform to lower 42 and 50 kDa adult isoforms that correlates with changes in cTNT splicing (23). Taken together, our results with Bru1 in *Drosophila* reflect a general requirement for CELF activity throughout myogenesis to regulate developmental transitions and fiber-type specific isoform expression.

Despite a requirement for Bru1 function throughout myogenesis, our data suggest that IFMs are sensitive to both the timing and dosage of Bru1 expression. Early overexpression of Bru1 with Him-Gal4 resulted in IFMs with a hole in the center of the myofiber, and rescue with Him-Gal4 enhanced myofiber, myofibril, sarcomere and contractile defects, indicating that too much Bru1 during early stages is detrimental to development (Fig 7). Early overexpression of CELF1 in mice with a β-actin or MCK promoter leads to central nuclei, reduced muscle mass, increased expression of p21, Myogenin and Mef2A, and lethality where phenotypic severity is directly correlated with the level of CELF1 overexpression (29,105). In culture, overexpression of CELF1 in myocytes promotes cell cycle and inhibits differentiation (106). Our data are consistent with and extend these results to show how overexpression of Bru1 affects myofibril assembly. Interestingly, recent work demonstrates that steady-state protein expression levels of CELF1 are significantly affected by use of alternative 3’-UTR regions (107), and that increased nuclear activity of CELF1 affecting alternative splicing but not increased cytoplasmic activity affecting translation in adult muscle leads to severe histopathology (108), revealing that muscles possess multiple mechanisms to finely-tune CELF expression levels. One mechanism that might explain Bru1 dosage effects is differential sensitivity of Bru1 targets. It has been shown that splicing of CELF1 targets Bin1 and Mef2A are only disrupted in a strong MHC-CELF/¢1 line and not in a milder line, indicating that different targets have different thresholds of responsiveness to CELF activity (88). Alternatively, dosage sensitivity may reflect interactions with other co-regulatory RBPs. RBPs are suggested to function in multi-factor complexes, such as the RBFOX1 containing LASR complex (109), where different combinations of constituent proteins lead to different regulatory outcomes. CELF proteins have been reported to have antagonistic interactions with both MBNL (21,24) and RBFOX family proteins (27,38). Around 22-30% of splice events are co-regulated by RBFOX2 and either CELF1 or CELF2 in mammalian heart, and RBFOX family motifs are enriched downstream of exons regulated after Bru1 RNAi in flies (27), suggesting functional conservation of antagonism. Further studies are needed to determine the molecular mechanisms underlying CELF family dosage sensitivity and to elucidate the co-regulatory logic of RBPs that participate in the fiber-type specific RNA-regulatory network.

Upregulation of CELF1 activity in differentiated muscle is thought to play a major role in the pathogenesis of myotonic dystrophy Type I (DM1) (OMIM 160900). CTG repeat expansions in the DMPK gene sequester and functionally deplete MBNL, leading to increased nuclear localization and activity of CELF1, with longer CTG repeats correlated with increased disease severity and decreased age of onset (110,111). CELF1 is normally downregulated 5-10 fold in adult vertebrate muscle (21,23,86) and reported to be required during early muscle development (88,100,112), such that increased nuclear activity in mature muscle results in a reversion to embryonic splicing patterns (21,25,28,30,108). In *Drosophila*, Bru1 levels also decrease as the IFMs develop (38), and our data show that while RNAi knockdown of *bru1* with late-stage Fln-Gal4 did not cause severe defects (Fig S6), overexpression of Bru1 impaired muscle function and sarcomere structure (Fig 8). When we rescued the *bru1^M3^* mutant with Fln-Gal4 driven UAS-Bru1, we were able to largely restore mature patterns of alternative splicing and partially-abrogate hypercontraction and myofiber loss, but we could not rescue myofibril and sarcomere structural defects (Fig 8). Given the parallels between Bru1 and CELF1 function in flies and vertebrates, our results provide mechanistic insight into the variable efficacy and phenotypes observed during the development of DM1 therapeutics (113,114). Current nucleic acid therapeutics or genome modification approaches are focused on restoring MBNL function by increasing MBNL expression levels, blocking MBNL binding to CTG-repeats or editing the DMPK locus (115,116), in turn decreasing CELF1 activity. Consistent with our rescue data in flies, clinical trials and drug studies in cells and mice often are able to restore mature patterns of alternative splicing and improve symptoms of myotonia (114,116,117), providing a positive outlook for the development of therapeutics that can significantly improve patient quality of life. However, our results indicate that a cure for DM1 will need to restore the balance in MBNL and CELF regulation to avoid dosage artifacts, and that timely intervention and early administration of therapeutics will be necessary to arrest the developmental progression that results in sarcomere structural defects and contractile deficits. Taken together, our results advance understanding of Bru1 function and more broadly highlight the importance of CELF protein dosage and the progressive nature of CELF phenotypes over time.

## Materials and Methods

### Fly stocks and crosses

Experimental work with *Drosophila melanogaster* was approved under German §15 GenTSV (license number 55.1-8791-14.1099). Fly stocks were maintained using standard culture conditions at room temperature. Fly food was made by combining 16 L water, 1,300 g corn flour, 150 g soy flour, 1,300 g molasses, 130 g agar, 300 g yeast, and 650 g malt extract in a water-jacketed cooker. After food was sufficiently cool, 415 mL of 10% Nipagin and 295 mL acid mix (3% phosphoric acid, 21% propionic acid) were added. Food was distributed in vials and bottles with a peristaltic pump, dried at room temperature and stored at 4 °C till use. All experimental crosses were kept in a 27 °C incubator.

*w^1118^* alone or in combination with the relevant Gal4 driver (Gal4 x *w^1118^*) was used as the wild-type control. The *bru1^M2^* and *bru1^M3^* alleles were generated by insertion of a selectable 3xP3-DsRed cassette using a CRISPR-Cas9 approach (118). The *bru1^M2^* mutant contains a cassette insertion upstream of *bru1* exon 12, and has been described previously (38). *bru1^M3^* is described in this manuscript. The UAS-Bru1 line was generated by amplifying the full-length *bru1-RA* transcript from *w^1118^* using RT-PCR (primer sequences are available in Table S4), cloning the cDNA into the *pUAST-attB* transformation vector containing a 5x UAS-hsp70 promoter region and a SV40 polyadenylation sequence, and integrating at the attP-86Fb landing site (119) with ýC31 integrase (Fig S8 A). RNAi against *bru1* (*bru1-IR*) was achieved with a previously characterized hairpin (GD41568) (14,38) obtained from the Vienna Drosophila Resource Center (VDRC). Df(2L)BSC407 is a deficiency allele on chromosome 2L that includes the *bru1* locus and was obtained from the Bloomington Drosophila Stock Center (BDSC). Mhc^10^ is a TDT and IFM-specific amorphic myosin mutant (120). Gal4 driver lines used in this study include: *Mef2*-Gal4 (121), which constitutively drives in all muscles; *Him*-Gal4 (37), which drives strongly in myoblasts and at early stages of IFM development until about 30 h APF; *salm*-Gal4 (80), which expresses in IFM starting from approximately 16 h APF; *Act88F*-Gal4 (78), which drives strongly in IFM starting at about 24 h APF; *UH3*-Gal4 (76), which expresses specifically in IFM starting at around 36 h APF; and *Fln*-Gal4 (79), which is expressed IFM-specifically from about 56 h APF (Fig S6 A). weeP26 is a GFP-trap line inserted in the intron between *Mhc* exon 36 and 37, which only tags the long isoform of *Mhc* that terminates in exon 37 after the insertion (39,61,122). Act88F localization was tracked using UASp-GFP-Act88F (72). GFP-tagged fosmid reporter fly lines included Strn-Mlck-GFP (strn^4^, which tags IFM-specific isoform R) (14), wupA-GFP (fTRG925), Kettin-GFP (fTRG477), Clip190-GFP (fTRG156), Mlp84B-GFP (fTRG678), and Mlp60A-GFP (fTRG709) (123). Talin-YPet is a CRISPR-mediated endogenous tag that strongly localizes to tendon attachment sites at muscle tips (124). All fly lines and reagents are listed in the Resources Table S5.

### Generation of the *bru1^M3^* CRISPR allele

*bru1^M3^* was generated by CRISPR-mediated genome modification, using a previously described approach (118). Two sgRNAs targeting sequences in the intron between *bru1* exon 17 and 18 and after the 3’-UTR downstream of *bru1* (Fig S1 A, Table S4) were screened in S2 cells for cutting efficacy and then co-injected with a 3x-P3 DsRed integration cassette flanked by approximately 1 kilobase long arms homologous to the genome sequence immediately upstream and downstream, respectively, of the sgRNA cut sites. The integration cassette contains a strong splice-acceptor followed by a three frame stop and a poly-adenylation sequence. Instead of the intended deletion, our genome modification resulted in the insertion of the cassette into the intron just upstream of exon 18, the last coding exon shared by all *bru1* isoforms (Fig S1A, B). Although we could detect increased expression of *bru1* mRNAs that include upstream exons 12 to 14, splicing into exon 18 was dramatically reduced, while splicing from exon 17 into the inserted construct containing the SV40 polyadenylation sequence was strongly increased (Fig S1 C). Splicing into the integrated cassette in *bru1^M3^* results in deletion of exon 18, which encodes the terminal 88 amino acids (aa516-604 of *bru1-RA*) in the third RRM domain (the extended RRM3 domain comprises aa471-604 (64)). *bru1^M3^* is therefore a truncation allele that results in early termination after exon 17, which affects all *bru1* isoforms (Fig S1 D).

### Behavioral assays

Flight ability was assayed as described previously (125). Briefly, N > 30 adult male flies were collected under CO_2_ on day 1 after eclosion, recovered overnight at 27 °C and introduced into a 1-meter-long cylinder divided into five zones. Flies with a “normal flight” ability landed in the top two zones, “weak fliers” in the middle two, and “flightless” males fell to the bottom. Eclosion competence was assayed by counting the number of adult flies that emerged from their pupal cases. At least 60 live pupae were unbiasedly selected after 48 h APF and monitored until eclosion.

### Immunofluorescence staining

Pupae of the desired genotype were tightly staged as follows: newly pupated 0 h APF flies (motionless, transparent-white body color and everted spiracles) were selected and sorted by sex, and males were transferred to a wetted filter paper in a 60 mm Petri dish and maintained at 27 °C until the desired timepoint. Pupae and adult flies were dissected and stained as described previously (126). For early pupae (before 48 h APF), we performed open-book dissections by removing the ventral half of the pupa to expose the developing IFMs. For late pupal (after 48 h APF) to adult fly stages, we bisected the thorax sagittally after fixation to allow visualization of IFMs. All samples were fixed in 4% PFA in 0.5% PBS-T (1× PBS + Triton X-100) for 30 to 60 minutes. For primary antibody staining, samples were blocked in 5% normal goat serum in 0.5 % PBS-T for 90 min at room temperature, and then incubated overnight at 4 °C with rabbit anti-Bru1 (1:500) (38) or rabbit anti-GFP (1:1000, Abcam ab290). All antibodies are listed in the Resources Table S5. After washing three times in 0.5% PBS-T for 10 minutes at room temperature, samples were incubated for 2 h at room temperature with secondary antibody Alexa 488 goat anti-rabbit IgG or rhodamine-phalloidin (1:500, Invitrogen, Molecular Probes). Samples were washed three times in 0.5% PBS-T and mounted in Vectashield containing DAPI.

### Cryosectioning and histological staining

Pupae were staged as above. Cryosections were performed as described previously (37). Samples were removed from the pupal case and fixed in 4% PFA in 0.5% PBS-T overnight at 4 °C, and then incubated overnight in 30% sucrose in 0.5% PBS-T at 4 °C on a rocking shaker. Pupae or 1 d adult thoraxes were arranged ventral side down in a vinyl specimen mold (Sakura Finteck), embedded in Tissue-Tek O.C.T. (Sakura Finteck) and snap-frozen on dry ice. Blocks were stored at -80 °C, and then sectioned from anterior to posterior at 30 µm on a cryostat (Leica). Sections were collected on glass slides coated with 0.44 mM chromium potassium sulfate dodecahydrate in 1% gelatin to avoid detachment of the tissue during subsequent washing steps. Slides were post-fixed for 5 min in 4% PFA in 0.5% PBS-T at room temperature, washed two times in 0.5% PBS-T, and stained with rhodamine-phalloidin (1:500) for two hours at room temperature in a humidity chamber protected from light. Slides were washed three times in 0.5% PBS-T, mounted with Fluoroshield containing DAPI (Sigma), and imaged on a Leica SP8X WLL upright confocal.

For hematoxylin-eosin staining, slides were post-fixed in formalin for 10 minutes, washed in 35 °C running tap water, and rinsed in distilled water. Slides were incubated with Harris Hematoxylin stain (Roth) for 1 minute, rinsed with distilled water, and placed under running tap water for 7 minutes to remove excess stain. After rinsing with distilled water, slides were stained in 1% Eosin solution (Apotheke Klinikum der Universität München) for 2 minutes, rinsed in distilled water and dehydrated in 70% EtOH, 96% EtOH, and 99% EtOH in consecutive 5-minute incubation steps. Slides were placed in xylenes (Roth) and mounted with Eukitt (Orsatec). After drying, slides were imaged on an Olympus IX83 inverted microscope with a 0.95 NA 40x objective.

### Confocal microscopy

Laser-scanning confocal images were acquired on a Leica SP8X WLL upright confocal using Leica LAS X software in the Core Facility Bioimaging at the LMU, Biomedical Center (Martinsried, DE). Sagittal hemi-thorax sections were imaged with an HCPL FLUOTAR 10x/0.30 objective to resolve the whole fiber morphology and with an HCPL APO 63x/1.4 OIL CS2 objective to capture myofibril and sarcomere structure. Fibers of early pupae (before 48 h APF) were imaged with an HCPL APO 20x/0.75 IMM CORR CS2 objective. Samples stained with rabbit anti-Bru1 were processed in replicates using same antibody mix and were imaged with the same laser gain settings. 3D stereo projection movies of GFP-Act88F and Mhc^Wee-P26^-GFP myofibril incorporation were assembled with Leica LAS X software from 0.1 mm confocal Z-stacks. Samples for the live imaging of spontaneous flight muscle contractions at 48 h and 72 h APF were prepared as previously described (127). 10-minute live recordings were obtained at a frame rate of 0.65 s / frame with an HCPL APO 40x/1.30 WATER CS2 objective. Samples for developmental GFP-tagged fosmid reporter expression at 48 h, 72 h and 1 d adult were stained with the same antibody mix and images acquired with the same laser settings on a Zeiss LSM 780 confocal microscope using a Plan-APOCHROMAT 100x/1.46 oil immersion objective lens.

### Transmission electron microscopy

*w^1118^* or *bru1^M3^* pupae were staged as described above to 48 h, 60 h, 72 h or 90 h APF, and removed from the pupal case. 70% of the abdomen was carefully removed in 0.1 M cacodylate buffer using fine scissors, and thoraxes were incubated in glutaraldehyde fix solution (4% glutaraldehyde in 0.1 M cacodylate solution with 3% sucrose) for 2 h at room temperature on a rocking shaker. After pre-fixation, pupae were cut sagittal with a #C35 Feather microtome blade (Feather), and then fixed overnight in glutaraldehyde fix solution on a rocker at 4°C. Samples were post-fixed in osmium tetroxide (2% in cacodylate buffer) and stained with uranyl acetate (2% in cacodylate buffer), dehydrated in a graded ethanol series, and embedded in epoxy resin following standard protocols. Samples were sectioned at 100 nm on an ultramicrotome (RMC MT 7000) with a diamond knife as previously described (128). Sections were collected on glass slides, stained with Richardson blue and evaluated on an Olympus BX61VS light microscope until the sectioning depth of the IFMs was reached. Sections were then collected on TEM grids, and post-stained with uranyl acetate and lead citrate to enhance contrast. Images were acquired on a 100 kV FEI Morgagni transmission electron microscope with a side-mounted SIS Megaview 1K CCD camera. At least two individuals per genotype and timepoint were analyzed.

### RNA isolation, RT-PCR and RT-qPCR

All primers are listed in Table S4. Whole thorax samples were prepared from 10 or more flies by removing the head, wings and abdomen in a drop of pre-cooled 1x PBS using a fine scissors. Dissected IFM (>30 flies) and TDT (>60 flies) muscle samples were prepared from 1 d adult flies as described previously (39). Legs were removed from >15 flies using fine scissors. Samples were snap-frozen in 50 μl of TRIzol (TRIzol Reagent; Ambion) on dry ice and stored at -80 °C. Total RNA was isolated using TRIzol-Chloroform per the manufacturer’s guidelines. RNA samples were treated with DNaseI (New England Biolabs) and concentration was assessed using a Qubit 2.0 Fluorometer (Invitrogen). For normalization purposes, equivalent quantities of total RNA were used for cDNA synthesis with the LunaScript RT SuperMix Kit (New England Biolabs). For RT-PCR, 1 μg of cDNA was amplified with Phusion polymerase for 32 cycles and separated on a standard 1% agarose gel together with a 100 bp or 1 kb ladder (New England Biolabs). Ribosomal protein L32 (RpL32, RP49) served as an internal control for some reactions. Semi-quantitative analysis of gel band intensity to determine differences in exon use was performed using the “gel analysis” feature in Fiji. Band intensity was normalized against RpL32 to compare expression levels, or was used to calculate percent exon usage as: 100 x (individual band intensity)/ Σ (intensity of all bands observed for same primer pair).

For RT-qPCR, total RNA was extracted from IFMs dissected from 150 flies using TRIzol. Reverse transcription and cDNA generation were performed with the LunaScript RT SuperMix kit, starting with 1 μg of cDNA. cDNA was diluted from 1:8 to 1:20, depending experimentally on the expression level of the target. Samples were assayed using SYBRgreen on a QuantStudio 3 (Applied Biosystems by Thermofisher Scientific) with an extension temperature of 60 °C for 40 cycles. Samples were normalized against either RpL32 or Vha44, and fold change was calculated using the 2^-ΛΛCT^ method (129). Data were plotted in GraphPad Prism, and significance was calculated from at least 3 replicates using either a student’s t-test (for a single sample versus control) or ANOVA (for more than two samples).

### Image Analysis

Image analysis was performed in Image J/Fiji (130). For every confocal-based assay, 10-15 images were acquired from >10 adult flies or pupae. For TEM, more then 20 images were acquired from 2-3 animals. DLM fiber integrity and length were measured from Z-stack projections of hemi-thorax samples based on rhodamine-phalloidin staining. Fiber length was measured as distance (μm) from anterior to posterior tip of a single fiber using the freehand drawing tool in Fiji. The same tool was used on TEM images, after setting the scale based on the inset in each image, to measure the sarcomere length and myofibril width as the distance between or across Z-disks, respectively. For confocal images, sarcomere length and myofibril width of sagittal sections, as well as myofibril diameter and density of cross-sections, were measured based on phalloidin (F-actin) staining using MyofibrilJ (37) (https://imagej.net/MyofibrilJ). The number of myofibrils per bundle was quantified in a semi-automated manner as follows: the boundaries of individual fiber bundles were determined from the Z-stack and at least 3 separate and complete bundles from > 10 flies were cropped, analyzed with MyofibrilJ, and then manually corrected for accuracy (myofibril “dots” were added or subtracted based on manual inspection of the MyofibrilJ output). Each “hook” or “ring” structured myofibril was counted as one. The number of myofibrils classified as “dots”, “hooks” or “rings” were manually counted per defined cross-sectional area. The number of actin cables per fiber (at 26 h and 28 h APF) was determined manually based on rhodamine-phalloidin staining. Bru1 signal intensity was analyzed as follows: the nuclei were selected based on DAPI staining using the wand tool in Fiji, and relative Bru1 signal in the nuclei (overlapping the DAPI positive regions) was quantified as the mean intensity of Bru1 staining divided by the mean intensity of the absolute background.

Quantification of spontaneous contractions in developing DLM has been described previously (37) and was modified here as follows. Live movies were obtained for 10 minutes, and “twitches” in each visible DLM fiber at 48 h APF were classified as single, double or triple contraction events, and resulted in displacement of the myofiber and return to the original resting position. The range of each single spontaneous contraction at 48 h APF was measured using the freehand drawing tool in Fiji as the distance between the tip of the fiber prior to and after the extension. At 72 hr APF, contraction events quantified in the *bru1* mutant were slower than “twitches” and resulted in a one-directional myofiber extension, i.e. without return to the original resting position. All data were tabulated in Excel, and plotting and statistical analysis were performed in GraphPad Prism 8.4.0, using ANOVA or unpaired Student’s t-test.

### Deconvolution

Deconvolution was performed in ImageJ using the Diffraction PSF 3D and FFTJ – DeconvolutionJ plugins. Images were obtained at a resolution of 23.2848 pixels per micron and a corresponding voxel size of 0.0429 x 0.0429 x 0.2014 micron^3^ for deconvolution and 3D projection. YZ stacks were generated with interpolation using the Reslice option native to ImageJ. A digital point spread function (PSF) was generated with the following settings: IR = 1.518, NA= 1.40, 580 nm, pixel spacing 23.28 and corresponding slice spacing of 4.97 units, with a Rayleigh resolution of 10.68 pixels. Deconvolution was performed without resizing at 32-bit, double precision, and with a gamma setting of 0.01/0.03.

### Proteomics

IFMs were dissected as described previously (39). 30 flies per genotype were dissected from *w^1118^* and *bru1^M2^* at 72h APF and 1 d adult, and 4 biological replicates were prepared per genotype. Samples were processed according to the manufacturer’s instructions using the PreOmics iST Sample Preparation Kit (Preomics, #0000.0061) and analysed by the Protein Analysis Unit (ZfP) at the LMU Biomedical Center as follows. Desalted peptides were injected in an Ultimate 3000 RSLCnano system (Thermo) and separated in a 25-cm analytical column (75µm ID, 1.6µm C18, IonOpticks) with a 100-minute gradient from 5 to 60% acetonitrile in 0.1% formic acid. The effluent from the HPLC was directly electrosprayed into a Qexactive HF (Thermo) operated in data dependent mode to automatically switch between full scan MS and MS/MS acquisition. Survey full scan MS spectra (from m/z 375–1600) were acquired with resolution R=60,000 at m/z 400 (AGC target of 3 x 10^6^). The 10 most intense peptide ions with charge states between 2 and 5 were sequentially isolated to a target value of 1 x 10^5^, and fragmented at 27% normalized collision energy. Typical mass spectrometric conditions were: spray voltage, 1.5 kV; no sheath and auxiliary gas flow; heated capillary temperature, 250 °C; ion selection threshold, 33,000 counts. MaxQuant 1.6.14.0 (131) was used to identify proteins and quantify by LFQ with the following parameters: Database, Uniprot_AUP000000803_Dmelanogaster_Isoforms_20210325.fasta; MS tol, 10 ppm; MS/MS tol, 20 ppm; Peptide FDR, 0.1; Protein FDR, 0.01 Min. peptide Length, 5; Variable modifications, Oxidation (M); Fixed modifications, Carbamidomethyl (C); Peptides for protein quantitation, razor and unique; Min. peptides, 1; Min. ratio count, 2. Additional analysis was performed in Perseus (132). We filtered the data to retain biologically relevant protein groups with missing intensities between mutant and control samples (MNAR values) by requiring at least three replicates in either control or mutant to contain a value, and we imputed missing values by replacement with a constant value (lowest observed intensity – 1). Differential expression was tested by t-test with FDR = 0.05. Results were exported and further visualization performed in R.

### mRNA-Seq and transcriptome bioinformatic analysis

For transcriptome analysis, IFMs were dissected from 1 d adult *w^1118^* and *bru1^M3^* flies as described previously (39). Two replicates of IFMs from 100 flies were dissected per genotype. RNA was isolated using Trizol and sent to LC Sciences (Houston, TX) for sequencing. After quality verification, poly-A mRNA selection and library construction, samples were sequenced as stranded, 150 bp paired-end on an Illumina HiSeq to a depth greater than 70 million reads. Transcriptome data from *bru1-IR* and control IFMs at 24 h, 30 h, 72 h and 1 d adult was generated previously (14,37), and was reanalysed as part of this manuscript.

Sequence data was mapped with STAR to ENSEMBL genome assembly BDGP6.22 (annotation dmel_r6.32 (FB2020_01)). Files were indexed with SAMtools and processed through featureCounts. Downstream analysis and visualization were performed in R using packages listed in the Table S5. Differential expression was analyzed at the gene level with DESeq2 and at the exon level with DEXSeq. Differential exon use can reflect alternative splicing as well as alternative promoter use. Both packages were additionally used to generate normalized counts values. We employed previously annotated sets of genes, including sarcomere proteins (37), genes with an RNAi phenotype in muscle (125), core fibrillar genes regulated by Spalt major and differentially expressed between IFM and tubular muscle (14), mitochondrial proteins (133,134), and all genes with the Flybase GO term “muscle contraction,” “actin cytoskeleton,” or “actin cytoskeleton organization”. Genes with a hypercontraction phenotype in *Drosophila* muscle were curated by hand from the literature. Full lists of all gene categories are available in Table S2. Plots were generated using ggplot2 or ComplexHeatmap.

### Data availability

Raw numbers used to generate plots are available in supplementary tables and raw data files. Images of all Western blots and RT-PCR gels are provided in the raw data files. mRNA-Seq data are publicly available from GEO with accession numbers GSE63707, GSE107247, GSE143430 and GSE205092. Whole proteome mass spectrometry data were submitted to Proteomexchange on June 22, 2023 and are available under accession number YYYYY.

## Acknowledgements

We sincerely thank Frank Schnorrer for advice, fly stocks and material support at the initiation of this project. MLS appreciated the generous support from Andreas Ladurner, as well as helpful discussions and shared facilities with Carla Margulies. We are grateful to Nicolas Gompel for access to injection equipment. We thank Sandra Lemke and Andi Pan for help setting-up the twitching assays, and Sabrina Chaabane for testing *bru1-IR* flight ability with a couple Gal4 drivers. We received generous access and support from Magdalena Götz and her department for cryosectioning. Peter Meinke and Stefan Hintze graciously helped with the H&E staining. We acknowledge receipt of fly stocks from the Bloomington Drosophila Stock Centre (BDSC) and the Vienna Drosophila Resource Center (VDRC). We acknowledge the Core Facility Bioimaging and the Protein Analysis Unit (ZfP) at the LMU Biomedical Center (Martinsried, DE) for mass spectrometry and confocal imaging support, respectively. We acknowledge the Deutsche Forschungsgemeinschaft (MLS, 417912216), the Deutsche Gesellschaft für Muskelkranke e.V. (MLS), start-up funding from the University of Missouri Kansas City (MLS), and the International Max Planck Research School (EN) for financial assistance.

## Author Contributions

Contributions are defined using CRediT role terminology (https://casrai.org/credit/).

Investigation (EN, MCG, AE, CG, JB, AR, HG, MH, MLS),

Writing – original draft (EN, MLS),

Writing – review & editing (EN, MLS, CB, MH, TS),

Conceptualization (MLS),

Data curation (TS),

Formal analysis (EN, MLS, TS, AR, MCG),

Visualization (EN, MLS, MCG, TS),

Resources (TS, HG, MH),

Supervision (MLS, MH, TS),

Funding acquisition (MLS, MH)

## Supplemental Figure Legends

**Fig S1. Molecular and phenotypic verification of the *bru1^M3^* CRISPR allele.**

**(A)** Diagram of the C-terminal region of the *bruno1* (*bru1*) locus and mRNA isoforms RA, RB and RD (exons, purple; UTRs, yellow). Location of the RNA recognition motif domains (RRM, light red), target region of anti-Bru1 antibody (brown), target region of *bru1-IR* GD41568 hairpin (brown), location of *bru1^M2^* construct insertion site (brown) and the sgRNAs (blue) used for CRISPR-mediated generation of the *bru1^M3^* hypomorph allele are marked. Transgenic construct is inserted upstream of exon 18 and contains a strong splice acceptor (SA, light blue), a triple frame stop (stop, red), an SV40 polyadenylation signal (orange) and a selectable 3xP3-dsRed marker (crimson) flanked by homology arms (light tan). Exon numbering according to the annotation FB2021-05. **(B)** Whole-fly genomic PCR verifying dsRed cassette insertion in the *bru1* locus. Identity of amplified region marked on the left, band size noted on the right. Primer sequences available in Table S4. **(C)** RT-PCR to test expression of *bru1* mRNA in whole-thorax and dissected IFM. Identity of amplified region marked on the left, band size noted on the right. RpL32 used as internal control. **(D)** Diagram of the *bru1^M3^* allele. Splicing from exon 17 is redirected into the splice acceptor of the inserted construct (red line, A), leading to early termination of the *bru1* mRNA and truncation of RRM3. Splicing from exon 17 to exon 18 is strongly reduced (dotted red line, B), and signal from 3’-UTR exon 21 is not detectable. **(E-F)** Confocal projections of 1d adult hemithoraxes showing IFMs from *bru1^M3^*/+ and *bru1^M3^*/Df(2L)BSC407. Deficiency BSC407 covers the complete *bru1* locus. Thorax boundaries in (F), dashed line; phalloidin stained actin, grey; Scale bar = 100 μm. **(E’-F’)** Single-plane confocal images of 1 d adult IFM myofibrils. Scale bar = 5 μm. **(G)** Quantification of myofiber phenotypes in (E-F). N > 40 myofibers/10 flies for each genotype. **(H-I)** Quantification of sarcomere length (H) and myofibril width (I) from TEM data shown in Fig 1 E. Boxplots are shown with Tukey whiskers, outlier data points marked as black dots. Significance determined by ANOVA and post-hoc Tukey (ns, not significant; ***, p-val < 0.001). **(J-K)** Quantification of Z-disc alignment (J) and sarcomere morphology (K) defects from TEM data shown in Fig 1 E. N > 20 images from 2 biological samples for each individual genotype and time point. **(L)** Dot plot showing the correlation between all detected peptide groups and their corresponding mRNA expression level in *bru1^-/-^*versus w^1118^ IFM (proteins with a significantly DE exon, orange; significantly DE genes, purple). The Pearson’s / Spearman’s correlation coefficients (top left corner) and regression line (blue) indicate a weak but positive correlation.

**Fig S2. Fiber-type specific alternative splicing of muscle proteins is disrupted in *bru1^-/-^* IFM.**

**(A)** Boxplot of gene (DESeq2), exon (DEXSeq) and protein level (mass spec) expression changes between 1 d adult *bru1^-/-^* and *w^1118^* IFM in select categories of genes including GO term “actin cytoskeleton organization”, microtubule associated genes, mitochondrial genes, GO term “muscle contraction”, RNA-binding proteins (RBPs), sarcomere proteins (SPs) and fibrillar core genes. Blue dot denotes p ≤ 0.05. **(B)** Venn diagram of the overlap between all significantly DE genes (purple), exons (orange) and proteins (green) between *bru1^-/-^* versus *w^1118^* IFM in 1 d adults. **(C-E)** RT-PCR verification of alternative splice events in *Strn-Mlck* (C), *wupA* (D) and *Mhc* (E). Top: scheme of alternative isoforms with primer locations. Exon numbering in accordance with the FB2021_05 annotation. Color coding of depicted isoforms consistent with bottom panel; 3’-UTR regions in light beige. Middle: Quantification of relative expression level of splice events in tubular leg and jump (tergal depressor of the trochanter, TDT) and fibrillar IFM, and in *bru1^M3^* IFM. Error bars = SD. Bottom: representative RT-PCR gel image. **(F-N’)** Misexpression of GFP-tagged sarcomere proteins in *bru1^M3^*IFM. (**F, I, L)** Diagrams of reporter GFP incorporation into tagged transcripts of *Strn-Mlck* (F), *wupA* (I), and *Mhc*-weeP26-GFP (L). Exons, magenta; 3’-UTR, tan; SA, splice acceptor; SD, splice donor; sGFP, superfold GFP. **(G-N)** Intensity matched single-plane confocal images from control and *bru1^M3^* IFMs at 90 h APF showing incorporation of Strn-Mlck-IsoR-GFP (G-H), wupA-GFP (J-K) and Mhc-weeP26-GFP (M-N). Strn-Mlck isoform R with sGFP tagged exon 25 is strongly expressed in wild-type IFM (G’) but absent from *bru1^M3^* (H’). WupA with an sGFP tagged exon 3 is normally absent from wild-type IFM (J’) but gained in *bru1^M3^*. Expression of the Mhc isoform containing exon 37 and tagged in weeP26-GFP is normally restricted to early IFM development (M-M’), but is altered in *bru1^M3^*(N-N’). GFP-tag, green; phalloidin stained actin, magenta (G, H, J, K, M, N); pseudo-colouring, GFP-intensity (compare G’ and H’; J’ and K’; M’ and N’). Scale bar = 5 μm. **(O-Q)** RT-PCR verification of alternative splice events in *Zasp52* (O), *Tm1* (P) and *sls* (Q). Top: schemes of alternative isoforms. Bottom: representative RT-PCR gel image. Labelled as in (C-E).

**Fig S3. Incorporation of both Act88F and Mhc into growing myofibrils is abnormal in *bru1^-/-^* IFM.**

**(A)** mRNA expression level raw C_T_ values assayed by RT-qPCR for different genes in *bru1^M3^* (red dots) and *w^1118^* (white dots) IFM, including *Act88F*, *Vps35*, *Lamp1*, *Vha44*, *mical*, *Nedd8*, *Atg2*, *Ald1*, and *His3.3B*. (two independent primer sets: N1 and N2, were used). Some genes, such as *His3.3B* and *Ald1* are changed in *bru1^M3^*, and not suited as a normalization standard. **(B-C)** Single plane confocal images of *Fln*-Gal4 driven pUAS-GFP-Actin88F incorporation into control (B-B’”) and *bru1^M3^* (C-C’”) at 90 h APF. Longitudinal sections (B, C) represent the XY-axis, while vertical lines mark the exact position of orthogonal slices at the z-disk (yellow line) and M-line (cyan line). Orthogonal view (B’-B’”, C’-C’”) represents YZ-axis of (B, C) respectively. GFP, green; phalloidin stained actin, magenta; Scale bar = 5 μm. **(D-G)** Single-plane confocal images of Mhc-weeP26-GFP expression in control (D-E””) and *bru1^M3^* (F-G””) at 90 h APF. Mhc-weeP26-GFP labels a specific isoform of Mhc that is only expressed during early IFM development. Longitudinal (D, F) and orthogonal sections (E-E””, G-G””) are shown as above at the z-disc (yellow line) and M-line (blue lines). GFP, green; phalloidin stained actin, magenta; Scale bar = 5 μm. **(H)** Violin plots of changes in expression of tubular-preferential genes and exons. Left plot shows changes in tubular-preferential gene and exon expression in tubular leg versus fibrillar wildtype IFM (yellow) and *bru1^M3^* versus wildtype IFM (red). Tubular-preferential was defined as all genes or exons with a log_2_FC > 1 and an adjusted p-value < 0.05 in the leg versus IFM comparison. Some but not the majority of tubular genes and exons are upregulated in *bru1^M3^* IFM. Right plot shows how the same tubular-preferential gene/exon sets change expression with time in IFM when comparing 1 d adult IFM to 24 h APF IFM in control (grey) or *bru1-IR* (orange) IFM. **(I)** Violin plots of changes in expression of fibrillar-preferential genes and exons. Left plot shows changes in fibrillar-preferential gene and exon expression in tubular leg versus fibrillar wildtype IFM (yellow) and *bru1^M3^*versus wildtype IFM (red). Fibrillar-preferential was defined as all genes or exons with a log_2_FC < -1 and an adjusted p-value < 0.05 in the leg versus IFM comparison, and thus expressed higher in IFM than in leg. Right plot shows how the same fibrillar-preferential gene/exon sets change expression with time in IFM when comparing 1 d adult IFM to 24 h APF IFM in control (grey) or *bru1-IR* (orange) IFM.

**Fig S4. Temporal dynamics of gene expression and exon use in *bru1-IR* IFM across muscle development.**

**(A)** Top: Boxplot of changes in gene expression across the *bru1-IR* timecourse (*bru1-IR* versus control) for temporal-switch genes that are normally upregulated (log_2_FC > 0, p-adj ≤ 0.05) or downregulated (log_2_FC < 0, p-adj ≤ 0.05) in control IFM from 24h to 1 d adult. Bottom: Boxplot of changes in exon use across the *bru1-IR* timecourse for temporal-switch exons that are normally upregulated (log_2_FC > 0, p-value ≤ 0.05) or downregulated (log_2_FC < 0, p-value ≤ 0.05) in control IFM from 24h to 1 d adult. Blue dot denotes p ≤ 0.05. **(B)** Boxplot of changes in gene expression (DESeq2) and exon use (DEXSeq) in GO term “actin cytoskeleton” genes in *bru1-IR* versus control IFM at 24 h, 30 h, 72 h APF and in 1 d adult. Blue dot denotes p ≤ 0.05. **(C)** Heatmap of gene level-expression changes in all sarcomere protein and fibrillar muscle genes at all timepoints in *bru1-IR* versus control IFM. The fifth column shows the temporal change in use of the same genes in wildtype IFM from 24h APF to 1 d adult. **(D)** Heatmap of all exons significantly DE (DEXSeq, p-val ≤ 0.05) at any timepoint in *bru1-IR* versus control IFM. The fifth column shows the temporal change in use of the same exons in wildtype IFM from 24h APF to 1 d adult.

**Fig S5. GFP-tagged sarcomere protein reporters reveal temporal misexpression dynamics in *bru1-IR* IFM.**

**(A)** Plot of mRNA-Seq based gene-level expression of *wupA* (maroon), *sls* (blue), *CLIP-190* (green), *Mlp60A* (purple) and *Mlp84B* (orange) in *bru1-IR* (dark colors) and control (light colors) IFM at 24h, 30h and 72h APF and in 1 d adult. log_2_(counts) of DESeq2 count values normalized across all four timepoints are plotted. **(B)** Plot of mRNA-Seq based exon-level expression of *wupA exon 1* (maroon), *sls exon 38* (blue), *CLIP-190 exon 29* (green), *Mlp60A exon 13* (purple) and *Mlp84B exon 2* (orange) in *bru1-IR* (dark colors) and control (light colors) IFM at 24h, 30h and 72h APF and in 1 d adult. log_2_(counts) of DEXSeq normalized count values are plotted. The selected exons contain the GFP-tag visualized in (C-G). **(C-E)** Expression of select splice-isoforms of *wupA* (C), *sls* (D) and *CLIP-190* (E) visualized by GFP-tag fluorescence (grayscale) in intensity-matched, single-plane confocal micrographs of IFM from control (left) and *bru1-IR* (right) flies at 48 h and 72 h APF and 1 d adult. The GFP-tag in *sls* labels the Kettin isoform. GFP, green; phalloidin stained actin, magenta; Scale bar = 5 μm. **(F-G)** Single plane confocal images of GFP-tagged Mlp60A and Mlp84B expression (greyscale) in control (left) and *bru1-IR* (right) IFM at 48 h and 72 h APF and 1 d adult. Expression is only detected in 1 d adult *bru1-IR* IFM. GFP, green; phalloidin stained actin, magenta; Scale bar = 5 μm.

**Fig S6. RNAi knockdown of *bru1* with temporally regulated Gal4 drivers reveals differential requirement for Bru1 during IFM development.**

**(A)** Scheme of temporal Gal4 expression during IFM myogenesis. All Gal4 drivers tested in this study are listed on the left and ordered by expression timepoint. Colored bars depict the time range when each Gal4 driver is expressed. Gal4 drivers used for RNAi and rescue experiments depicted in main Figs have a distinct color (*Him*-Gal4, tan; *UH3*-Gal4, turquoise; *Fln*-Gal4, yellow). Gradient color of the bar indicates the strength of temporal expression. Key time-points in IFM myogenesis are marked at the bottom. **(B)** Quantification of the percent of flies that eclose from pupal cases in Gal4 controls and *bru1-IR*. No eclosion defect was noted for any of the *bru1-IR* lines tested. **(C)** Quantification of flight ability in Gal4 controls and *bru1-IR* knockdown flies. N > 45 flies for each genotype. **(D)** Quantification of myofiber ripping and detachment phenotypes in Gal4 controls and *bru1-IR* knockdown at 90 h APF and 1d adult flies. N > 40 fiber for each genotype and time-point. **(E-P)** Confocal projections of hemi-thoraxes showing DLMs of *salm*-Gal4, *Act88F*-Gal4 and *Fln*-Gal4 driven *bru1-IR* at 90 h APF (E-J) and 1 d adult (K-P). The myofibers of *salm*-Gal4 and *Act88F*-Gal4 driven *bru1-IR* are already ripped at 90 h APF (F, H), while *Fln*-Gal4 driven *bru1-IR* myofibers remain intact (J, P). Scale bar = 100 μm. **(E’-P’)** Single-plane confocal images of genotypes as in (E-P) showing myofibril and sarcomere phenotype of *bru1-IR* at 90 h APF (E’-J’) and 1 d adult (K’-P’). Scale bar = 5 μm. **(Q-R)** Quantification of sarcomere length (Q) and myofibril width (R) in (E’-P’). Boxplots are shown with Tukey whiskers, outlier data points marked as black dots. Significance determined across the time-course by ANOVA and post hoc Tukey (ns, not significant; \*\**P* < 0.01; \*\*\**P* < 0.001).

**Fig S7. Validation of *bru1* RNAi knockdown efficiency with different temporal Gal4 drivers at the mRNA and protein level.**

**(A)** RT-qPCR verification of *bru1* gene expression levels in mutant and knockdown conditions in 1 d adult IFM. Expression is shown relative to the matched control, either wildtype *w^1118^* or a Gal4 driver crossed to *w^1118^*. *bru1* levels were strongly and significantly reduced in *bru1^M3^*, *bru1-IR*, *bru1-IR^Him^*, *bru1-IR^UH3^* and *bru1-IR^Fln^*. **(B-G)** Single-plane confocal images of IFM nuclei stained with rabbit anti-Bru1 in control, *bru1^M3^*, *bru1-IR^UH3^* and *bru1-IR^Fln^* at 90 h APF. Bru1 signal is absent in *bru1^M3^* (C-C’) and *bru1-IR^UH3^*(E-E’) IFMs, but can still be detected in *bru1-IR^Fln^*IFMs (G-G’). Images were acquired using same settings and pseudo-coloured based on intensity (B’-G’). Bru1, green; DAPI, magenta; Scale bar = 5 μm. **(H)** Quantification of Bru1 relative signal intensity based on fluorescence levels in (B-G). Boxplots are shown with Tukey whiskers, outlier data points marked as black dots. Significance determined by ANOVA and post hoc Tukey in comparison to matched control (ns, not significant; \*\*\**P* < 0.001). **(I-P)** Single-plane confocal images of IFM nuclei stained with rabbit anti-Bru1 in control and *bru1-IR^Him^*at 24 h, 30 h, 48 h and 90 h APF. Bru1signal is absent from *bru1-IR^Him^* IFM at 24 h (M-M’) and 30 h (N-N’) APF, but can be detected at 48 h (O-O’) and 90 h (P-P’) APF. Images were acquired using same settings and pseudo-coloured based on intensity (I’-P’). Bru1, green; DAPI, magenta; Scale bar = 5 μm. **(Q)** Quantification of Bru1 relative signal intensity based on fluorescence levels in (I-P). Data visualized as in (H). Significance determined by ANOVA and post hoc Tukey in comparison to matched control (ns, not significant; \*\**P* < 0.01; \*\*\**P* < 0.001)

**Fig S8. Overexpression of Bru1 pre- or post-myofibrillogenesis generates strong fiber and myofibril phenotypes.**

**(A)** Diagram of the UAS-Bru1-RA (UAS-Bru1) expression construct integrated into the attP-86Fb landing site on chromosome 3R. The construct contains a 5x UAS-hsp70 promoter region, full-length *bru1-RA* coding sequence and an SV40 terminator and polyadenylation sequence. **(B)** Quantification of the percent of flies that eclosed from control and UAS-Bru1 overexpression with Him-Gal4, salm-Gal4, Act88F-Gal4, UH3-Gal4 and Fln-Gal4. Overexpression with salm-Gal4 is embryonic lethal. **(C)** Quantification of flight ability in control and UAS-Bru1 overexpression with Act88F-Gal4 and UH3-Gal4. N > 30 flies for each genotype. **(D)** Quantification of myofiber ripping and detachment phenotypes at 90 h APF and 1 d adult in control and UAS-Bru1 overexpression with Act88F-Gal4 and UH3-Gal4. N > 40 fibers for each genotype and time-point. **(E-N)** Confocal projections of hemi-thoraxes from control and Act88F-Gal4 and UH3-Gal4 driven UAS-Bru1 at 90 h (E-I) and 1 d adult (J-N). Myofibers of Act88F-Gal4 Bru1 overexpression are fully degraded (G, L). Dashed line outlines the thorax boundaries in (G, L). Scale bar = 100 μm. **(E’-N’)** Single-plane confocal images showing myofibril and sarcomere phenotypes at 90 h APF (E’-I’) and 1 d adult (J’-N’). Scale bar = 5 μm. **(O-P)** Quantification of sarcomere length (O) and myofibril width (P) in (E’-N’). Boxplots are shown with Tukey whiskers, outlier data points marked as black dots. Significance determined for each time-point by ANOVA and post hoc Tukey (ns, not significant; \**P* < 0.05; \*\**P* < 0.01; \*\*\**P* < 0.001).

**Fig S9.Expression of Bru1 with *Him*-Gal4 is early stage-specific and sufficient to generate phenotypes in control and *bru1^M3^* IFM.**

**(A)** Single-plane confocal images of IFM nuclei stained with rabbit anti-Bru1 in Him-Gal4 overexpression of UAS-Bru1 at 24 h, 48 h and 90 h APF. Bru1 signal is detected at 24 h APF, but not at 48 h or 90 h APF. Images were acquired using same settings and pseudo-coloured based on intensity. Bru1, green; DAPI, magenta; Scale bar = 5 μm. **(B)** Quantification of Bru1 relative signal intensity based on fluorescence levels in (A). Horizontal line denotes the mean value of Bru1 signal intensity in *bru1^M3^*. **(C)** Confocal projections of hemithorax and single plane images of IFM myofibrils at 48 h and 90 h APF and 1 d adult in control, *Him*-Gal4 driving UAS-Bru1, and *Him*-Gal4 driving UAS-Bru1 in a heterozygous mutant background (*bru1^M3/+^*). Phalloidin stained actin, grey; Scale bar = 100 μm (hemithorax), or 5 μm (myofibrils). **(D)** Quantification of DLM fiber integrity at 90 h APF. Genotypes denoted by symbols: top row, *bru1* allele presence (wild-type *bru1^+/+^*, white square; heterozygous *bru1^+/-^*, half-red square, mutant *bru1^M3-/-^*, red square); middle row, *Him*-Gal4 driver presence (absent, empty; present, tan triangle); bottom row, UAS-Bru1 presence (absent, empty; present, magenta rhombus). N > 40 fibers for each genotype. **(E-F)** Quantification of sarcomere length (E) and myofibril width (F) in (C). Boxplots are shown with Tukey whiskers, outlier data points marked as black dots. Significance determined for each time-point by ANOVA and post hoc Tukey (ns, not significant; \**P* < 0.05; \*\*\**P* < 0.001). **(G)** Histological stain with hematoxylin and eosin (H&E) in wild-type, mutant and Him-Gal4 rescue IFM myofibers at 48 h APF. Hole, yellow arrowheads; Scale bar = 100 μm. **(H)** Quantification of myofiber morphology in (G). N > 10 for each genotype. **(I)** Quantification of myofibril width in (Fig.7 Q-T”). Data plotted and significance is determined as in (E-F).

**Fig S10. Expression of Bru1 with Fln-Gal4 produces mild sarcomere defects in control but restores alternative splicing defects in *bru1^M3^* IFM.**

**(A)** Single-plane confocal images of IFM nuclei stained with rabbit anti-Bru1 in control and rescue at 90 h APF. Images were acquired using same settings and pseudo-coloured based on intensity. Bru1, green; DAPI, magenta; Scale bar = 5 μm. **(B)** Quantification of Bru1 relative signal intensity based on fluorescence levels in (A). Statistical significance determined by unpaired t-test (\**P* < 0.05). **(C)** Confocal projections of hemithorax and single plane images of myofibrils at 90 h APF and in adult in control, *Fln*-Gal4 driving UAS-Bru1, and *Fln*-Gal4 driving UAS-Bru1 in a heterozygous mutant background (*bru1^M3/+^*). Phalloidin stained actin, grey; Scale bar = 100 μm (hemithorax), or 5 μm (myofibrils). **(D-E)** Quantification of sarcomere length (D) and myofibril width (E) in (C). Genotypes marked by symbols as in Fig S9. Boxplots are shown with Tukey whiskers, outlier data points marked as black dots. Significance determined for each time-point by ANOVA and post hoc Tukey (ns, not significant; \*\*\**P* < 0.001). **(F)** Quantification of myofibril width in (Fig 8 R) at 90 h APF. Significant determined as in (D-E). **(G-H)** RT-PCR verification of alternative splice events in *Zasp52* (G) and *Mhc* (H). Top: scheme of alternative isoforms with primer locations. Exon numbering in accordance with FB2021_05 annotation. Color coding of depicted isoforms consistent across top, middle and bottom panels; 3’ UTR regions in light beige. Middle: Quantification of relative expression level of detectable events in control *w^1118^* leg, jump (tergal depressor of the trochanter, TDT) and fibrillar IFM muscle, as well as mutant and Fln-Gal4 rescue IFM. Error bars = SD. Bottom: representative RT-PCR gel image. **(I)** RT-PCR verification of alternative splice events in *Tm1*. Top: scheme of alternative isoforms. Bottom: representative RT-PCR gel image. Splice events in *Tm1* were detected with distinct reverse primers, as isoforms do not share a common 3’-UTR.

**Table S1.** File

**Table S2**

**Table S3.**

**Table S4.** Primer table. Complete list of all primer sequences used in this manuscript.

**Table S5.** Key resources table. Complete list of all antibodies, fly lines, software packages, etc. used in this manuscript and their source.

