## Supplementary figures and images for "Bruno 1 regulates cytoskeleton dynamics and a temporal splicing transition to promote myofibril assembly, growth and maturation in *Drosophila* flight muscle"

### FigureS1

Nikonova et al, Figure S1

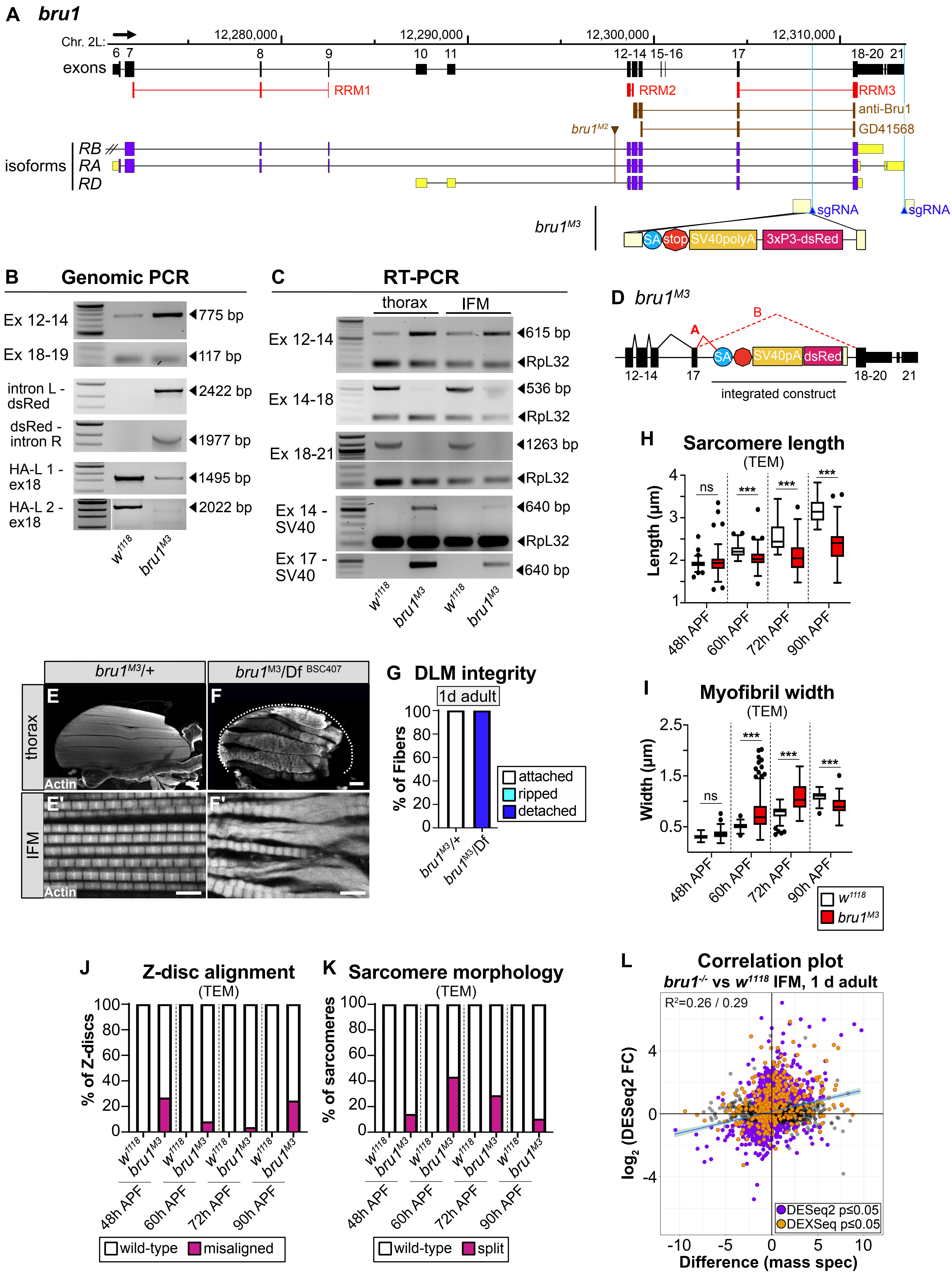

### FigureS2

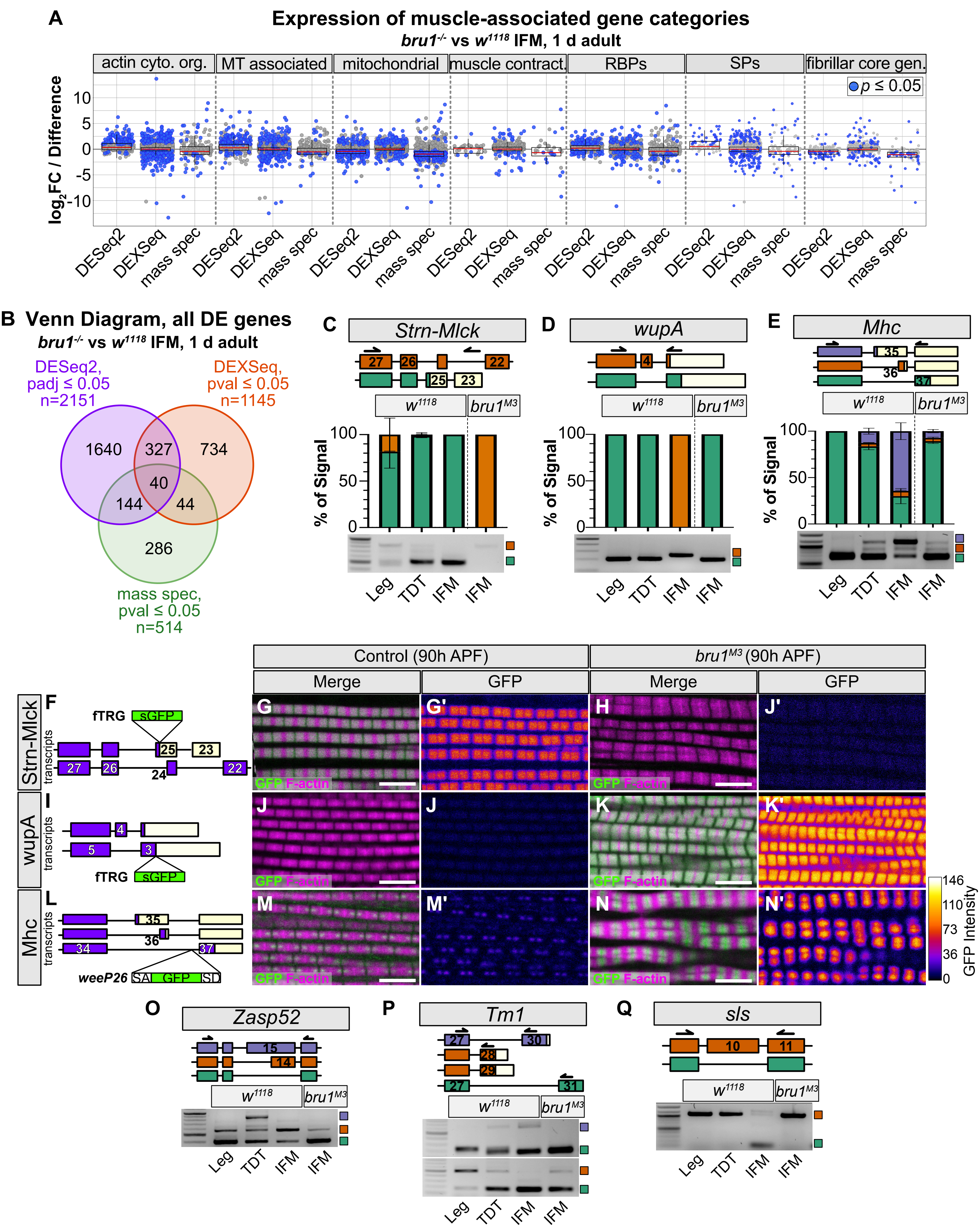

### FigureS3

**Nikonova et al, Figure S3**

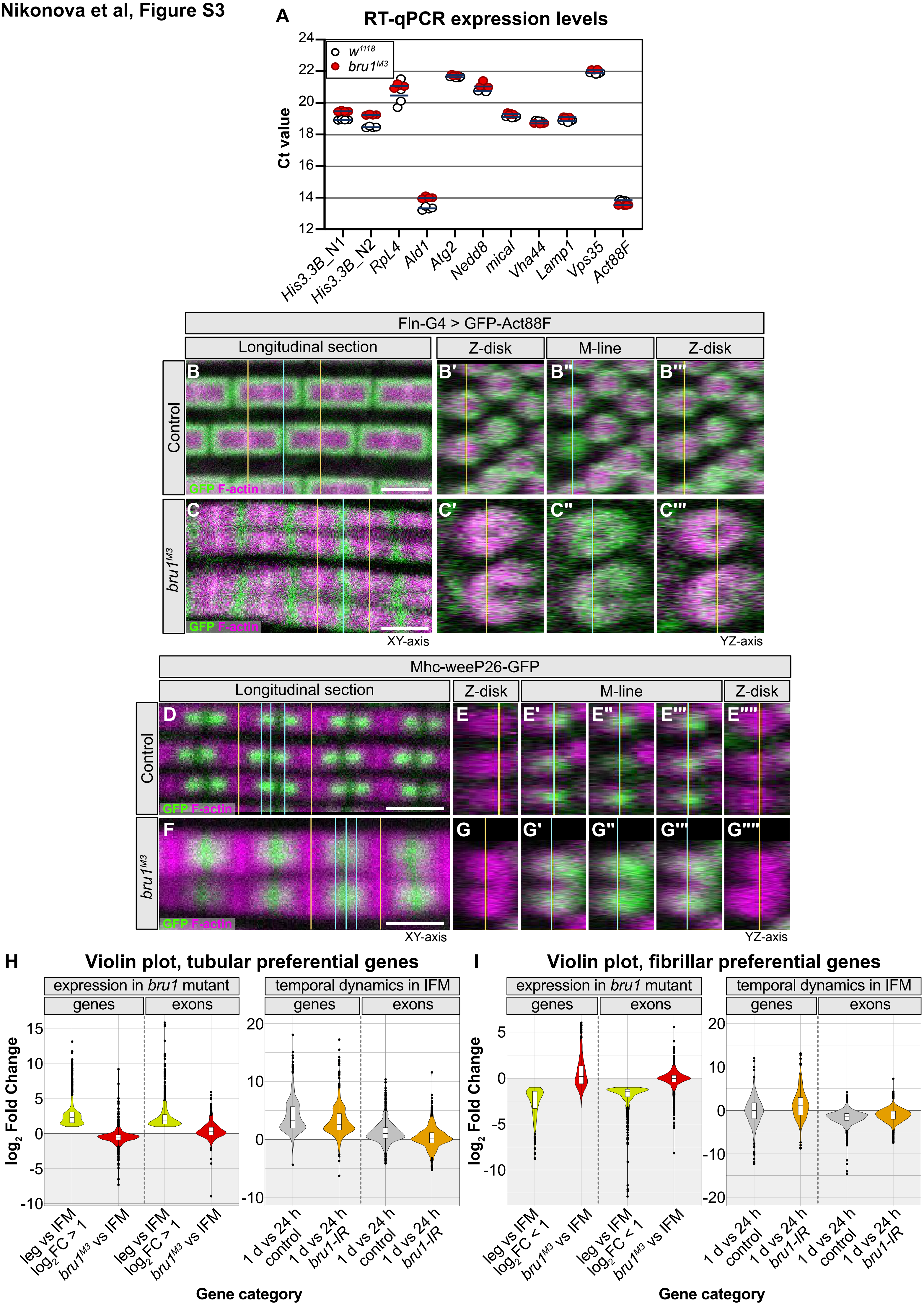

### FigureS4

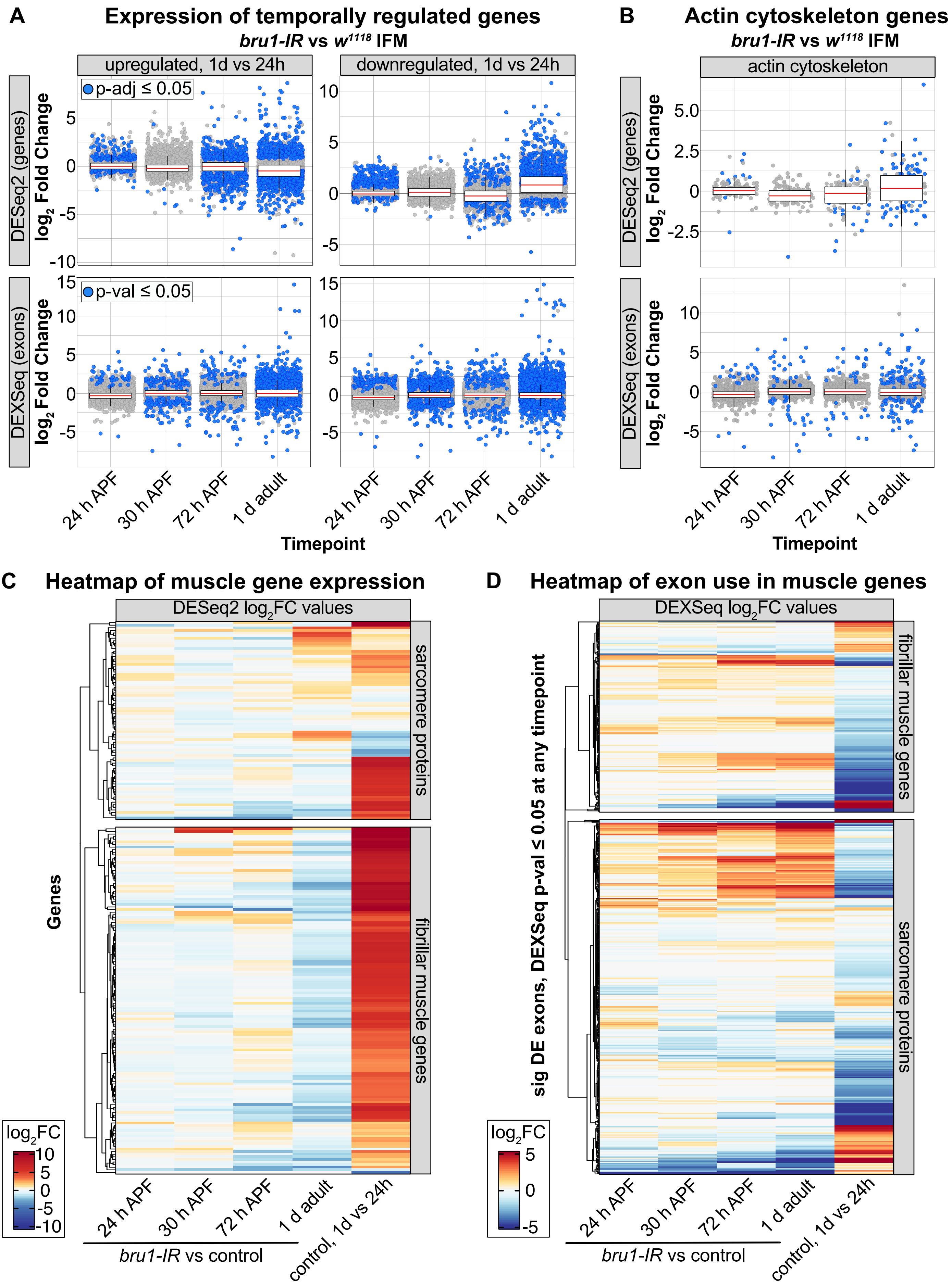

### FigureS5

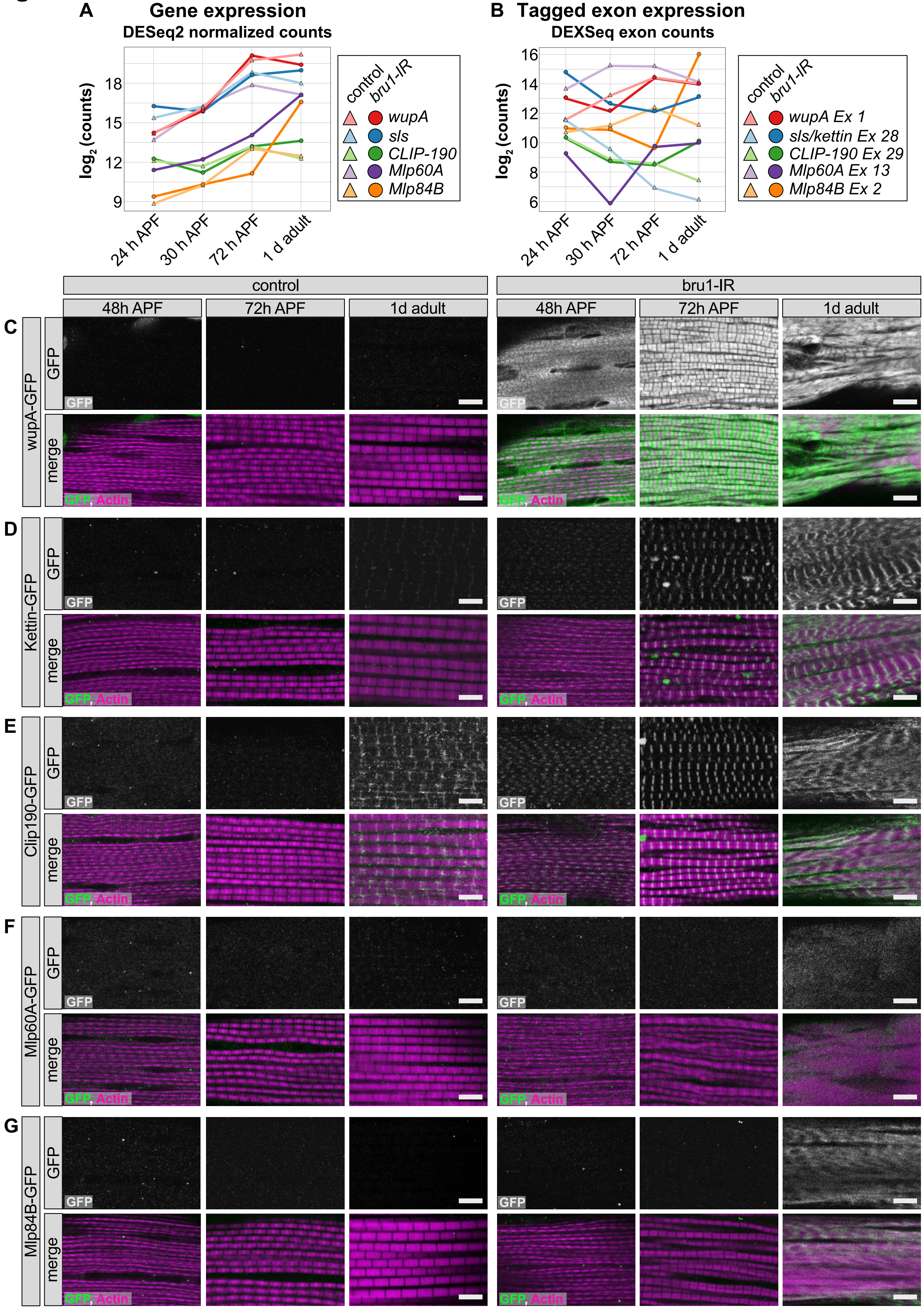

### FigureS6

Nikonova et al, Figure S6

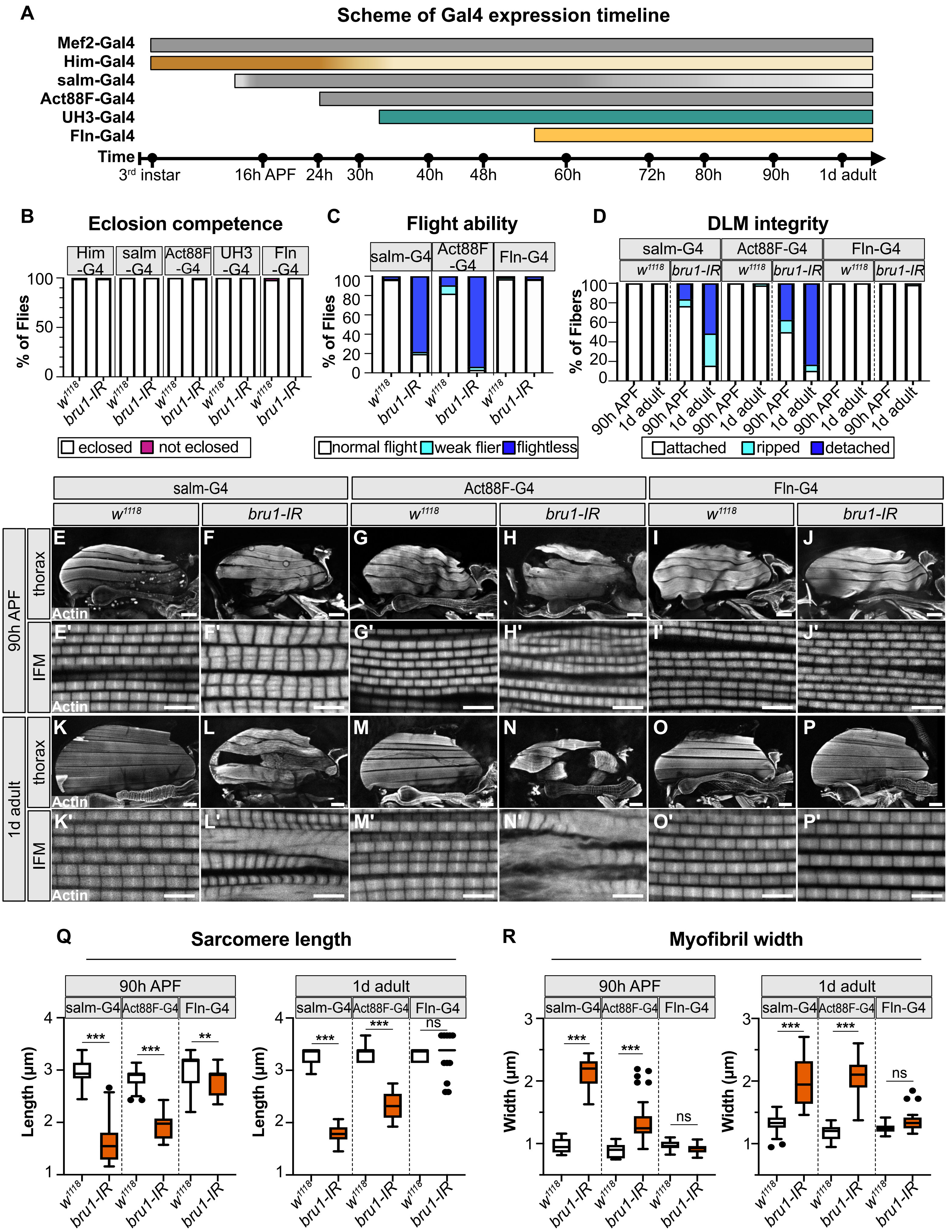

### FigureS7

Nikonova et al, Figure S7

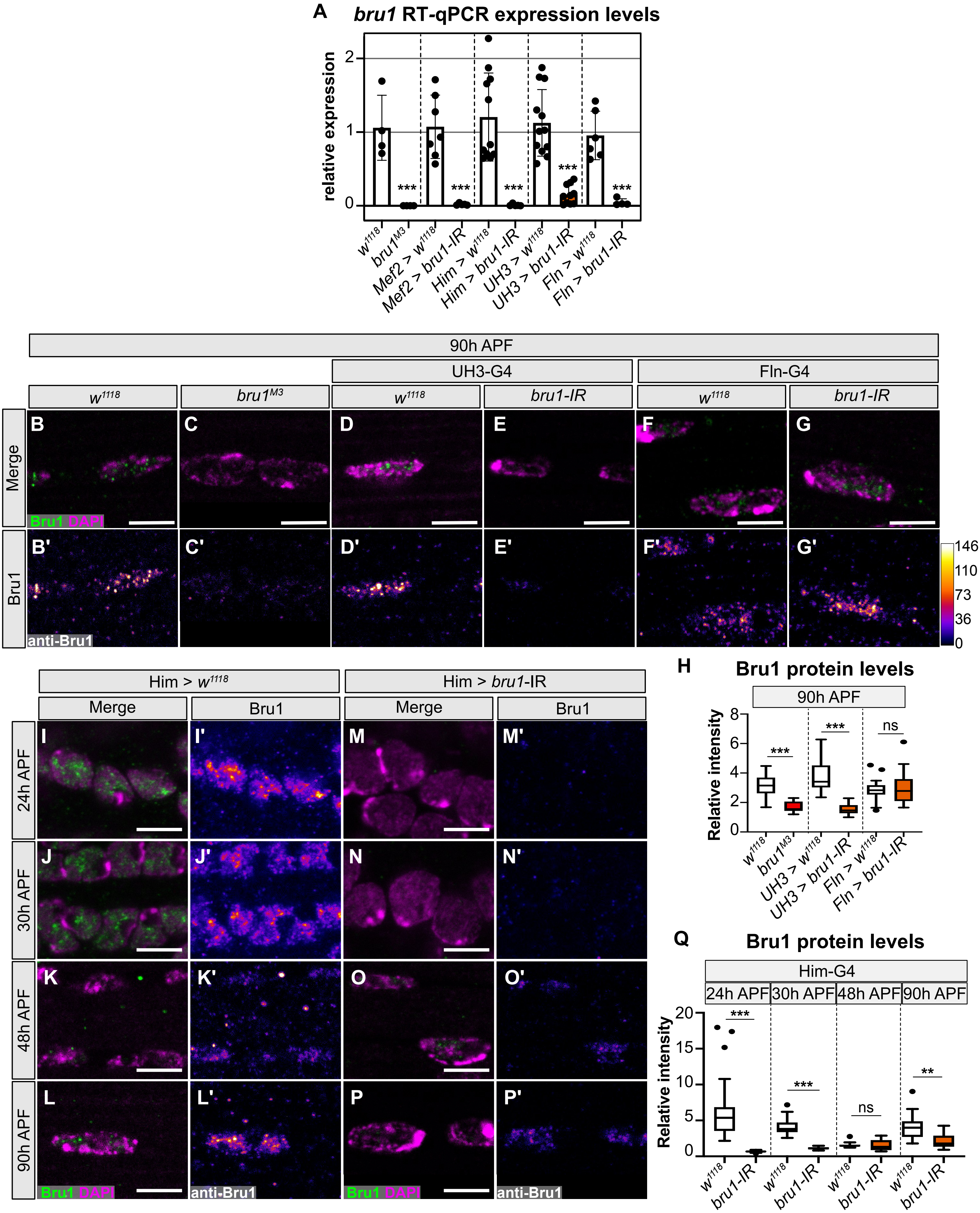

### FigureS8

Nikonova et al, Figure S8

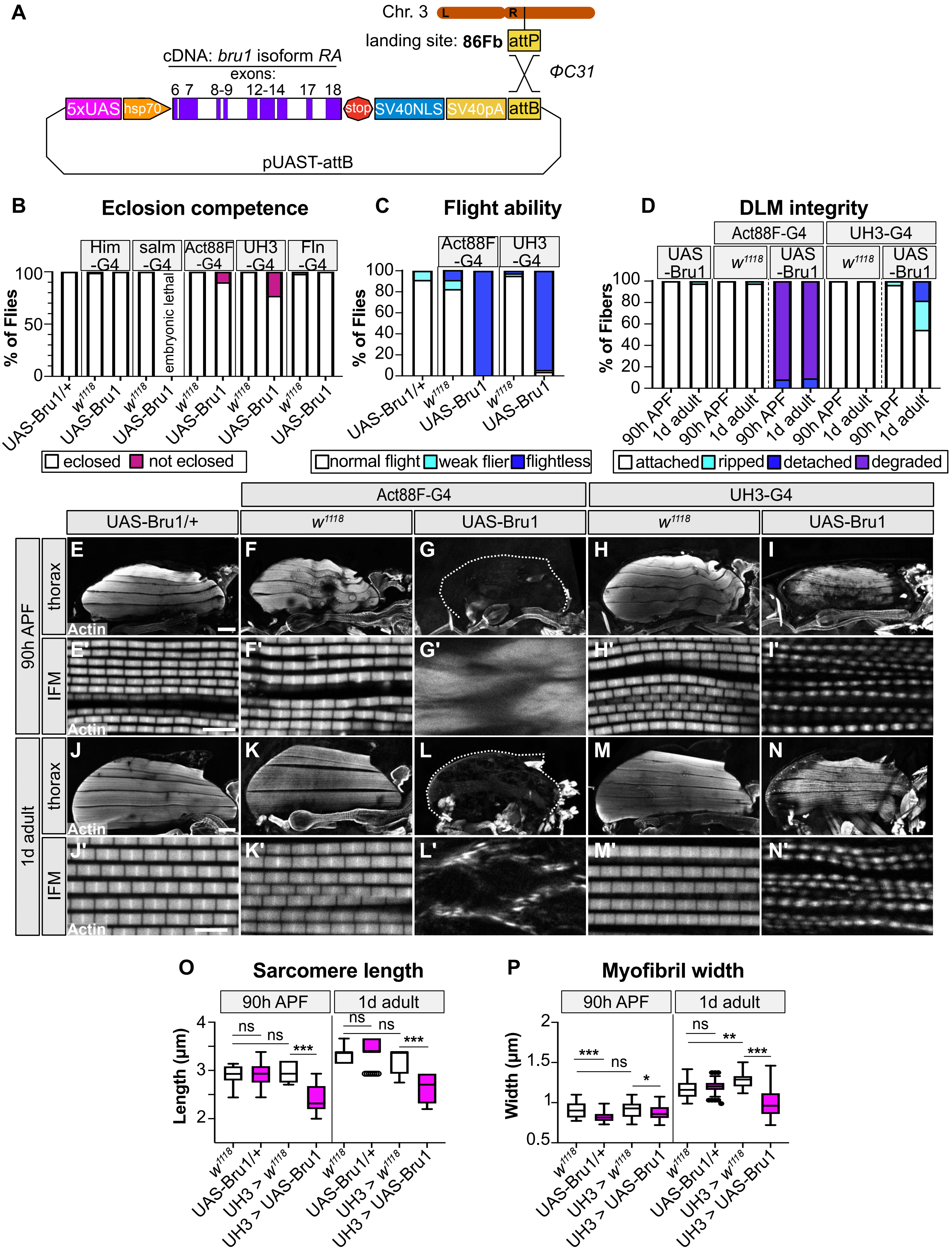

### FigureS9

Nikonova et al, Figure S9

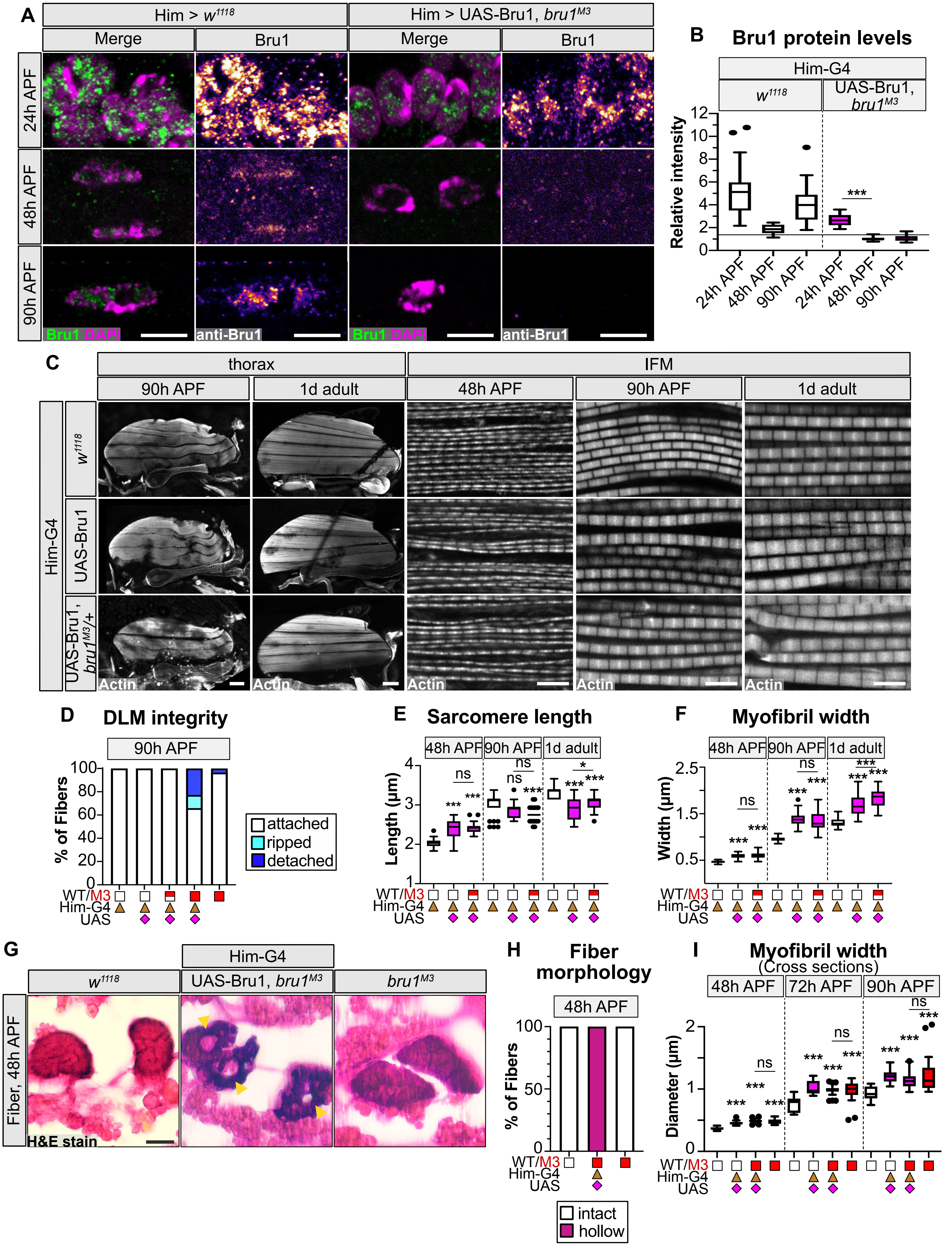

### FigureS10

**Nikonova et al, Figure S10**

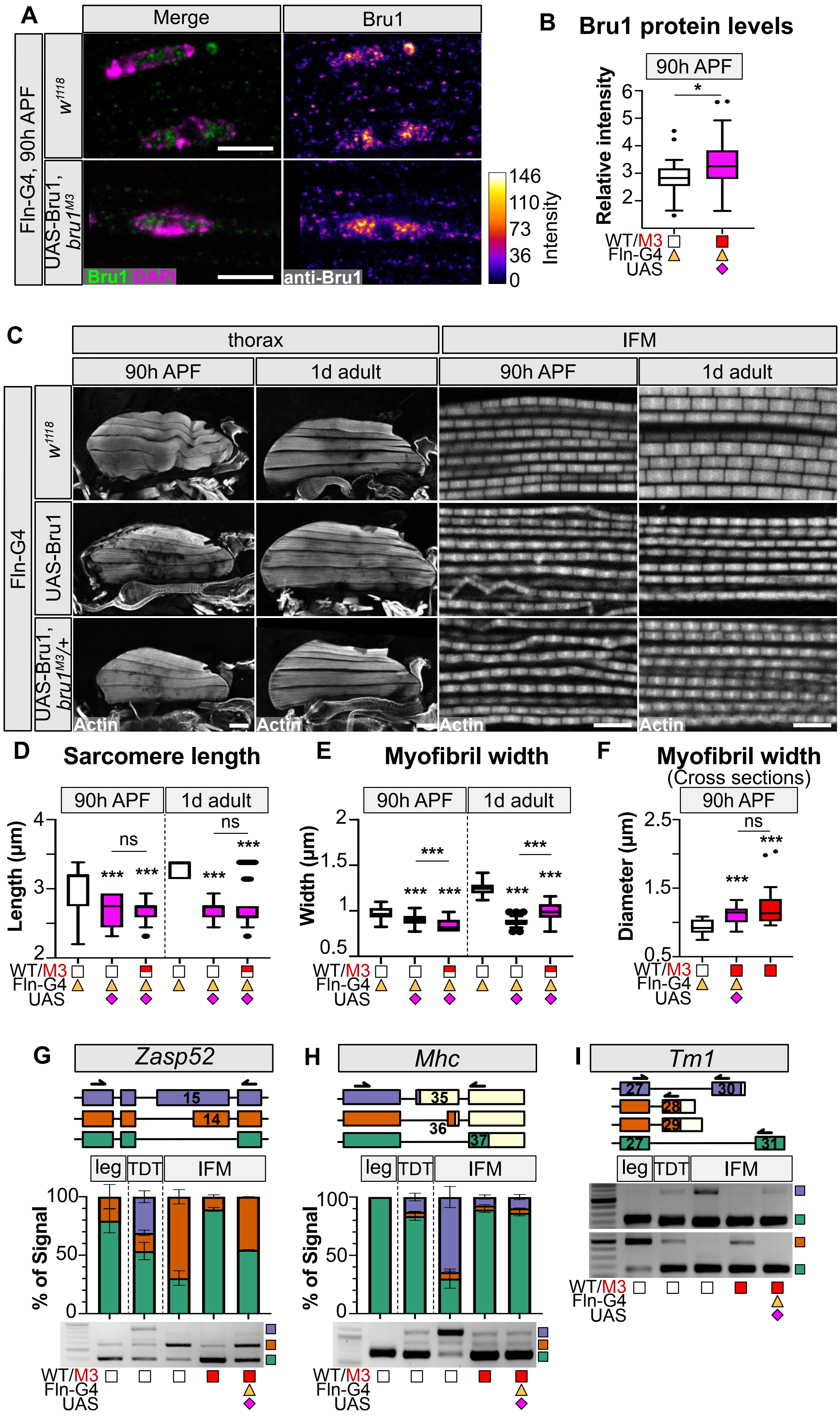
